# Single-phage profiling illuminates viral individuality during cell fate determination

**DOI:** 10.64898/2026.02.20.707030

**Authors:** Ehsan Homaee, Wenqing Zhu, Tianyou Yao, Ido Golding

## Abstract

The choice between cell death (lysis) and viral dormancy (lysogeny) following bacteriophage infection serves as a founding paradigm for the emergence of cellular heterogeneity in a genetically uniform population. The determination of host fate arises through the stochastic transcription from multiple viral genomes present within each cell, but this activity remains hidden from empirical interrogation, which typically stops at the whole-cell level. Here we use parallel sequential fluorescence in situ hybridization (par-seqFISH), followed by spatial clustering of phage-encoded transcripts within each cell, to profile the transcriptional activity of individual phages during synchronized infection of *Escherichia coli* (*E. coli*) by bacteriophage lambda. At the whole-cell level, transcription kinetics capture the developmental choice between lysis and lysogeny, and further demonstrate that viral replication is required for the emergence of diverging fate decisions. Zooming in to the single-phage level illuminates an individuality of viral activity during infection. We find that, while cells pursuing lysogeny display consensus activity of all inhabiting phages, lytic cells may contain phages that exhibit lysogenic activity. These findings support an earlier suggestion that consensus among coinfecting phages is required for cell dormancy. More broadly, our results highlight the need to identify how whole-cell behavior emerges from the activity of physically distinct copies of the same genetic circuit.

## INTRODUCTION

Bacteriophage infection served as an early paradigm for the emergence of phenotypic heterogeneity among genetically identical cells [1, 2]. In particular, the choice between lysis and lysogeny following infection of *E. coli* by phage lambda became the founding example for binary cell fate choice [3, 4] and for the purported role of biochemical stochasticity in driving diverging outcomes within a uniform population of cells [5–7]. During this process, the fate of the infected cell emerges from the stochastic transcription of multiple identical phage genomes that coexist within the same cell, due to both coinfection [8, 9] and ongoing viral replication [10, 11]. However, single-phage activity remains hidden under current empirical methods, where this information is coarse-grained to the whole-cell level [11–13].

Here, we seek to overcome this limitation and illuminate the transcriptional activity of individual phages during lambda infection and the subsequent choice between lysis and lysogeny. To achieve this goal, we first utilize par-seqFISH [14–16] to quantify the expression of phage genes in individual cells at successive time points following synchronized infection at a controlled multiplicity of infection (MOI) [11] (Figure 1A). The single-cell data exhibits a clear divergence of cell fates into the lytic and lysogenic attractors, at the expected population fractions [11]. Consistent with an earlier hypothesis [11], we find that active viral replication is required for the emergence of diverging developmental trajectories.

**Figure 1.**
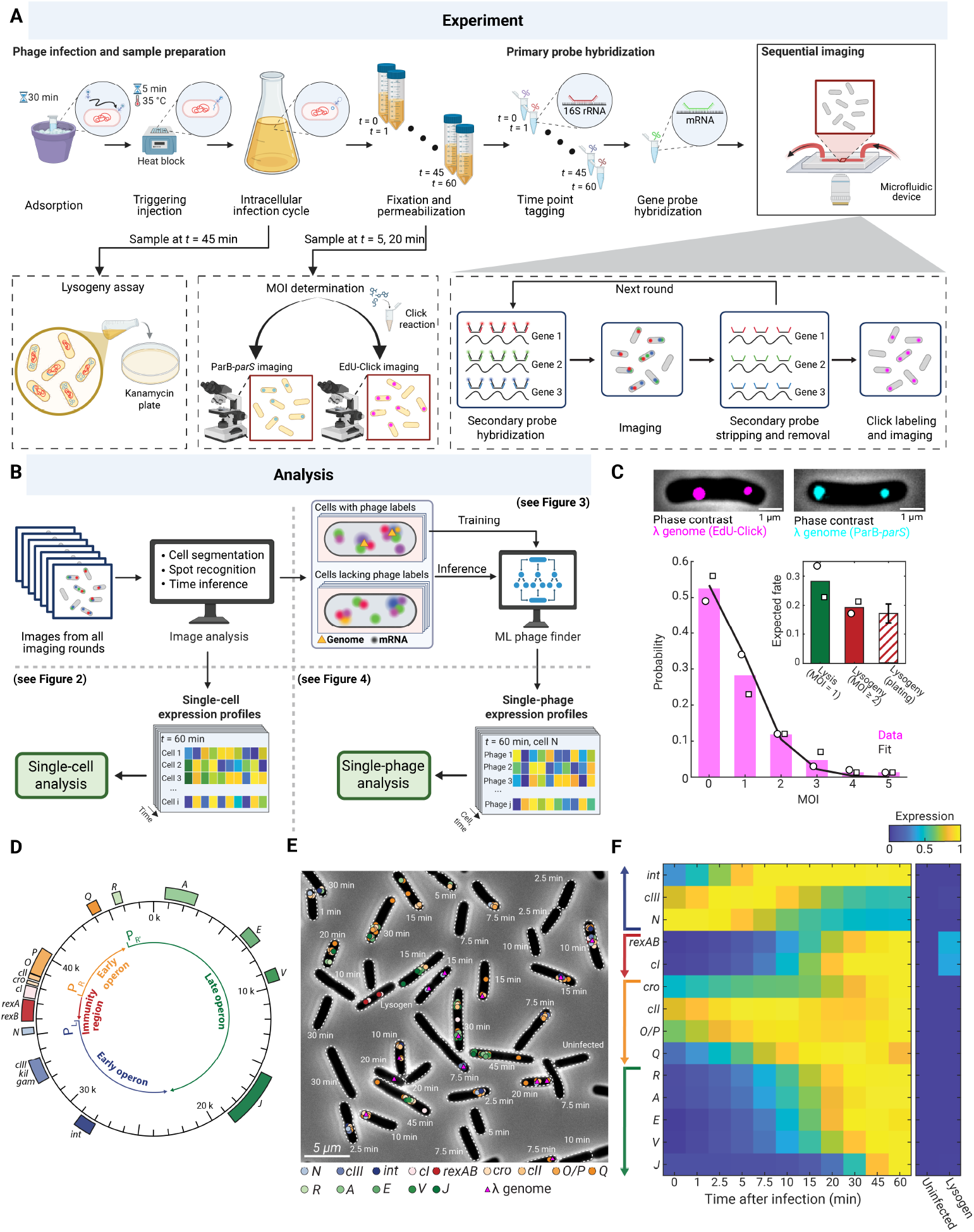
Par-seqFISH of synchronously infected cells captures the temporal program of phage transcription. (A) **Experimental workflow**. Phage adsorption is followed by a step of triggering genome injection, then initiation of the intracellular infection cycle. At the indicated times after infection, cells are harvested, fixed, and permeabilized. A subset of these samples are used for MOI determination using genome (ParB-*parS* and EdU-Click) imaging. In addition, cells are collected and plated on LB agar plates supplemented with kanamycin, to estimate the frequency of lysogenization. The time point of each fixed and permeabilized sample is barcoded using probe sets targeting the 16S rRNA. The samples from different times are then pooled and hybridized using phage gene probes. Cells are loaded into a microfluidic device for sequential imaging cycles, each consisting of secondary probe hybridization, imaging, secondary probe stripping and removal, and, finally, Click reaction and imaging of EdU-labeled phage DNA. Created with BioRender.com. (B) **Analysis workflow**. An image-analysis pipeline is used to perform cell segmentation, spot recognition, and time inference. These outputs are processed to yield the expression profiles of individual cells, used for single-cell analysis (Figure 2). Separately, spatial analysis of RNA spots and phage DNA labels provides training data for a machine-learning-based model that infers phage positions in the absence of phage genome label (Figure 3). The inferred phage positions, together with RNA information, define single-phage expression profiles within each cell, which are used for single-phage analysis (Figure 4). Created with BioRender.com. (C) **Measuring the multiplicity of infection in individual cells**. Top, *E. coli* cells with the infecting phages labeled using EdU-Click (left) and ParB-*parS* (right). Cells were imaged 20 min after infection. Bottom, the distribution of multiplicity of infection (MOI) in individual cells, measured by EdU-Click labeling. Circles and squares indicate data from *t* = 5 min (*n* = 145 cells) and *t* = 20 min (*n* = 175 cells), respectively. Black line, fit to a Poisson distribution (with an inferred mean of 0.63). The corresponding distribution measured using ParB-*parS* is shown in Figure S1. Inset, the fraction of cells expected to undergo each fate, based on the measured distribution of single-cell MOI. Cells with MOI = 1 are expected to become lytic (green bar), whereas cells with MOI ≥ 2 are predicted to become lysogens (red bar; both estimated from the two time points). The true fraction of lysogens was measured by plating on antibiotic plates (dashed bar; error bars indicate SE from bootstrapping; see METHODS). (D) **Bacteriophage genes targeted in this study**. The genetic map of *λ* is shown, with the measured transcripts colored according to their promoter. (E) **Multiplexed RNA detection and time-point annotation in individual cells**. A representative field of view is shown, with segmented cells, detected transcripts (color coded as in panel Figure 1D; each spot may represent more than one mRNA copy) and EdU-Click-labeled phage genomes (purple triangles). Cells were annotated according to their sample time point, or as belonging to the uninfected and lysogen control samples. (F) **Expression kinetics of** *λ* **following infection**. The cell-averaged mRNA copy number for each gene, as a function of time after infection (*n* = 31 – 160 infected cells per time point) was smoothed by fitting to a sum of two Hill functions, and the resulting values were normalized by the maximal value across all samples. Arrows on the left indicate the corresponding phage operons. Also shown (right) are the expression profiles in lysogen (*n* = 91 cells) and uninfected (*n* = 251 cells) controls.

Next, we leverage the colocalization of nascent (actively transcribed) mRNA molecules [17] to infer the spatial positions of individual phage genomes within each cell (Figure 1B). Inference is performed by training a machine-learning model, using fluorescently labeled phage genomes to provide the ground truth. This procedure allows us to profile the transcriptional activity of coinfecting phages during the choice of cell fate. We find that cells pursuing lysogeny display lysogenic transcription activity by all phages present. In contrast, lytic cells may contain phages of both lytic and lysogenic profiles. This finding suggests the dominance of the lytic choice, resulting in the need for intracellular consensus to enable dormancy [9].

## RESULTS

### Par-seqFISH of synchronously infected cells captures the temporal program of phage transcription

We first aimed to capture the expression kinetics of phage genes in individual cells undergoing lysis and lysogeny. To do so, we performed synchronized infection of *E. coli* bacteria (strain MG1655) using bacteriophage lambda (*λ cI857 Sam7 stf* ::P1*parS-kan*^R^) [11] (see METHODS, Table S1). To ensure that both lytic and lysogenic outcomes are represented in the population, we aimed for an average intracellular MOI of approximately 1 [18], and confirmed this value by single-cell microscopy of the infected cells, utilizing fluorescent labeling of the incoming phage genomes [11, 19] (Figure 1C, Figure S1; see METHODS). Given this average multiplicity, the near-Poissonian distribution of single-cell MOI [9, 11] results in ∼20% of cells being infected by two or more phages (Figure 1C), and thus likely to undergo lysogeny [11]. The fraction of lysogenic cells was confirmed by plating on selective antibiotic plates (Figure 1C).

We collected samples of infected cells at different times following phage entry. As controls, we used an uninfected culture and a lysogenic strain (MG1655 *λ cI857 stf* ::P1*parS-kan*^R^). The individual samples were barcoded using probes targeting 16S rRNA, following the method of [15, 16]. The pooled sample was then labeled and imaged using the seqFISH workflow of [15, 16] (Figure 1A). To capture the progression of phage development, we chose to target 18 genes (Figure 1D) representing the key lambda promoters [20, 21]. For especially long transcripts and for key fate-determining factors, we targeted more than one gene per polycistronic transcript (Figure 1D). Following imaging, the original samples were demultiplexed following [15, 16], and mRNA copy numbers for each gene inferred in individual cells following [22] (Figure 1E).

The expression kinetics of lambda genes are shown in Figure 1F, Figure S2, and Figure S3. The synchronized infection procedure allows the phage transcription program to clearly unfold, beginning with the “early” (*cro, N*) and “delayed early” genes (*cIII, cII, O, P, Q*) from the P_L_ and P_R_ promoters. Consistent with the expectation that both fate choices will be represented in the infected population, we observed the expression of both lytic (*R, A, E, V, J*, from P_R’_) and lysogenic (*cI, rexAB*, from the immunity region) genes, at the expected temporal order [23](Figure 1F, Figure S2, and Figure S3). Moreover, the progression of transcription along the lambda genome is consistent with the elongation speed of RNAP [17, 24] (Figure S4). The measured mRNA values were highly correlated across independent biological repeats (Figure S2, Figure S5, Figure S6), as well as with RNA-seq data obtained using the same infection protocol [25] (Figure S5; see METHODS). Taken together, the data indicates that applying par-seqFISH to synchronously infected cells faithfully captures the temporal program of phage transcription.

### Single-cell profiling reveals diverging developmental choices and demonstrates the requirement for viral replication

As noted above, our expectation from the distribution of single-cell MOI (Figure 1C) was that both lytic and lysogenic cells will be represented in the population. The presence of both lytic and lysogenic transcripts in the population-averaged expression (Figure 1F) supported this premise. We next sought to delineate individual cells pursuing the two routes.

To do so, we performed principal component analysis (PCA) of the single-cell RNA data [26] (Figure 2A; see METHODS). The first two principal components (PC1, PC2) together captured > 60% of the total variance (Figure S7), and their coefficients (loadings) were highly similar across biological repeats (Figure S8). Moreover, the vectors lent themselves to simple interpretation (Figure 2A): The first principal component (PC1) distinguishes active viral reproduction—indicated by transcription of early or lytic genes—from dormancy, where only the lysogeny maintenance gene *cI* (and the co-transcribed *rexAB*) are present. Thus, a cell’s position along this axis indicates whether it is pursuing lysis (positive values) or lysogeny (negative values). This interpretation is consistent with the bimodal distribution of PC1 values observed at late time points (Figure 2B). The second principal component (PC2) is captures developmental time, by weighting phage genes according to the temporal progression of transcription from the viral genome [21, 23] (compare Figure 2A with Figure 1F). Consistent with this interpretation, the cell-averaged value of PC2 increases with the sampled time after infection (Figure 2B).

**Figure 2.**
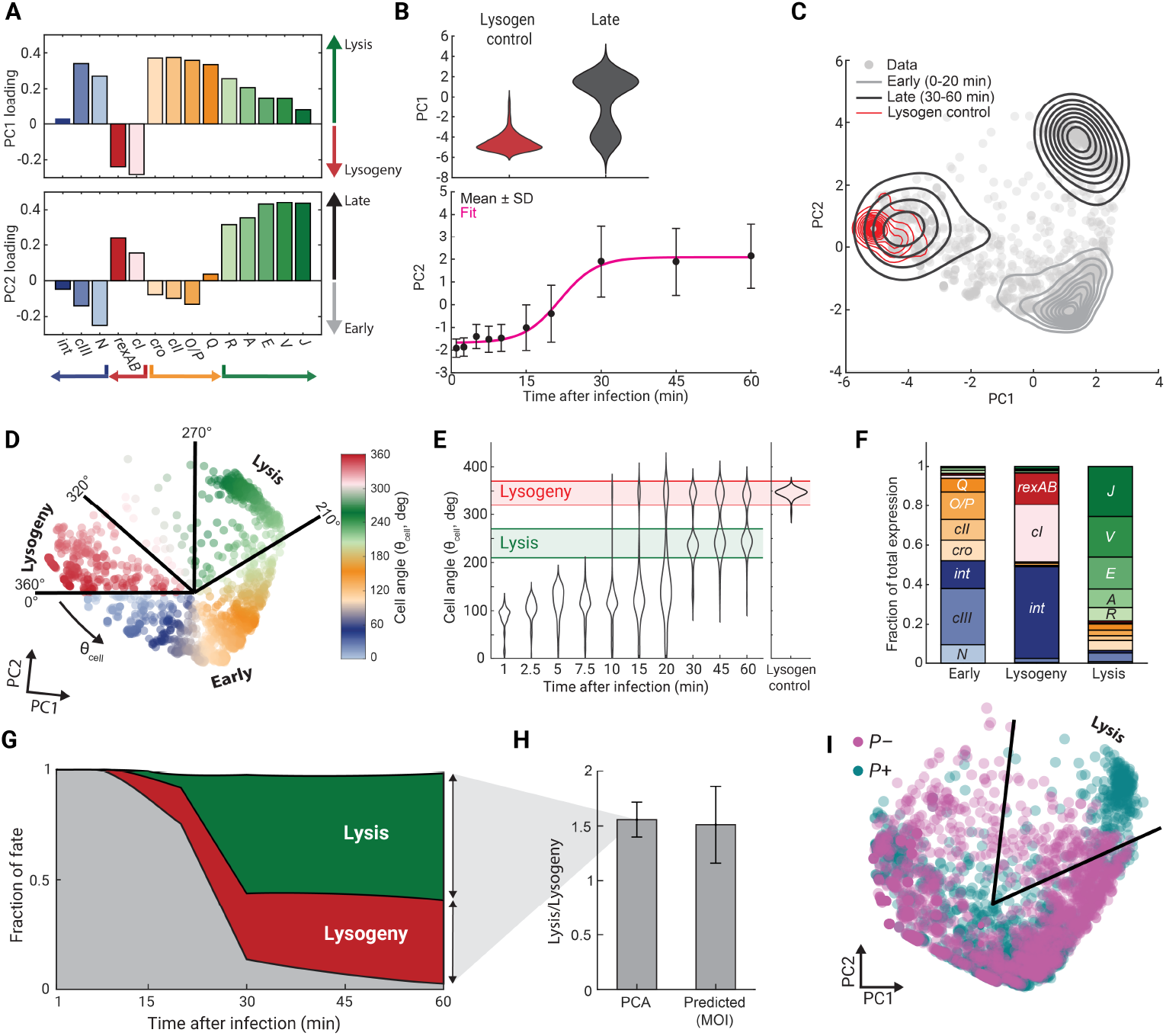
Single-cell profiling reveals diverging developmental choices and demonstrates the requirement for viral replication. (A) **PCA eigenvectors of the single-cell gene expression data**. The coefficents (loadings) of the first two principal component vectors derived from the single-cell RNA dataset (*n* = 1,768 cells). *λ* genes and operons are colored as in Figure 1D. (B) **Interpretation of the eigenvectors**. Top, PC1 scores for lysogen control (*n* = 91 cells) and for late-stage infected cells (pooled from *t* = 30, 45, 60 min, *n* = 221), with the latter exhibiting both lytic (PC1>0) and lysogenic (PC1<0) populations. Bottom, PC2 scores as a function of time after infection (*n* = 31 – 160 cells per sample), demonstrating the monotonic increase with time. Black markers, mean ± SD at each time point; pink line, fit to a sigmoid function, serving as a guide to the eye. (C) **Infected cell states in the PC1–PC2 plane**. The expression profile of each cell was projected to the plane of PC1 and PC2. Gray points, individual cells; solid lines, kernel-density contours for early-stage (0–20 min, light gray, *n* = 889), late stage (30–60 min, dark gray, *n* = 221), and lysogenic control cells (red, *n* = 91). (D) **Mapping cellular state to the polar cell angle** *θ*_*cell*_. Individual cells are shown in the PC1–PC2 plane, colored by the value of *θ*_*cell*_. Radial boundaries define sectors corresponding to early, lytic, and lysogenic regions, as indicated. (E) **The distribution of cell angles as a function of time after infection**. Violin plots show the distribution of *θ*_*cell*_ at each time point (*n* = 31 – 160 cells per sample) and for the lysogenic control sample (*n* = 91). Colored regions correspond to the classification of cell fates defined in panel D. (F) **Expression profiles of** *λ* **genes in the different developmental regions**. The expression of each target gene was summed over all cells within each region (defined using the *θ*_*cell*_ values as in panel D), and converted to the fraction of the total transcript count across all genes and cells in the region. (G) **The emergence of cell fates during infection**. Based on the angular regions defined in panel D, cells at each time point were scored as lytic (green), lysogenic (red), or early (gray). The colored areas indicate the fraction of cells in each category over time. (H) **The inferred fraction of cell fates agrees with MOI-based prediction**. Left bar, the ratio of cell numbers classified as lytic and lysogenic at 30-60 min after infection; error bar indicates the SEM across the three time points. Right bar, the ratio of cell numbers with MOI = 1 (predicted to lyse) and MOI ≥ 2 (predicted to form lysogens), inferred from the click-based MOI measurements at 5 and 20 min; error bar indicates the SEM across the two time points. (I) **Non-replicating phages fail to exhibit lytic expression**. PCA was performed on the combined single-cell datasets from *P*+ (teal, *n* = 1,768 cells) and *P*– (magenta, *n* = 2,203 cells) infections. The resulting PC1–PC2 scores are shown for individual cells.

The biological interpretability of the two principal components above suggests that the developmental trajectory of infected cells can be naturally mapped to the plane of PC1 and PC2 (Figure 2C). In this representation, temporal progression is captured by motion “up” from negative to positive PC2, while the choice of cell fate manifests in movement to either the left (lysogeny, PC1 < 0) or right (lysis, PC1 > 0). This simple picture is supported by the data (Figure 2C): Early-infection cells (*t* = 0 – 20 minutes) are centered in the lower-half plane (PC2 < 0) near the horizontal center line (PC1 = 0), whereas late infected cells (*t* = 30 – 60 minutes) form two separate clusters in the upper-half plane, one on each side of the PC1 = 0 line. The left (PC1 < 0) cluster largely overlaps the position of the control population consisting of a lysogenic strain, as expected for infected cells undergoing lysogenization. The right (PC1 > 0) cluster consists of cells undergoing lytic development, an inference supported by the transcription profile of cells within this cluster (see below).

The correspondence between infected cells’ developmental trajectories and position in the PC1–PC2 plane motivated us to simplify the representation further by describing the transcription state of each cell using the polar angle of its position, which we denote as the “cell angle” (*θ*_cell_) [27] (Figure 2D). As *θ*_cell_ = 0, we choose the polar direction that maximizes the discrimination between early-infected and lysogenic cells [28, 29]. The inferred value of this direction (≈ 6^◦^ below the negative PC1 axis) is consistent with our interpretation of the principal components (Figure 2A above). In the new representation, early infection, lytic, and lysogenic profiles are mapped, respectively, to *θ*_cell_ values of 0^◦^ – 210^◦^, 210^◦^ – 270^◦^, and 320^◦^ – 360^◦^ (Figures 2D–2F and Figure S9). This simplified one-dimensional description facilitates tracking the temporal evolution of cell state, by examining the single-cell values of *θ*_cell_ during infection (Figure 2E). Doing so is illuminating: Cell states center around the “early” attractor up to ≈ 15 min post infection, when the population diverges into the lytic and lysogenic attractors. The ratio of cells following the lytic and lysogenic routes, inferred from this analysis (≈ 1.6, Figure 2G), is consistent with the expected value based on the measured distribution of single-cell MOIs (Figure 2H), further supporting the validity of the analysis.

In earlier work, utilizing a replication-deficient phage mutant (*λ cI857 Pam80 stf* ::P1*parS-kan*^R^, denoted *P*–), we failed to detect a clear MOI-dependent divergence of cell fates [11]. Using a mathematical model of the decision circuit, we inferred that the absence of genome replication prevented phages infecting at low multiplicity from reaching the critical levels of Cro and Q to allow lytic commitment, instead arriving at a transient “failed lytic” state [11]. To revisit this idea, we performed par-seqFISH during infection by *P*– phage (Figure S3, Figure S6, Figure S10). Applying PCA to the single-cell data resulted in principal vectors similar to those of the wild type (*P*+) phage (Figure S11), consistent with the identicality of the decision circuit in the two strains. The positions of cells in the developmental plain (PC1–PC2), however, exhibited a marked difference: *P*– infected cells in the lytic quadrant remained closer to the origin than *P*+ infected cells, consistent with a scenario of failed lysis due to insufficient expression of lytic-driving genes (Figure 2I). The lysogenic attractor, in contrast, was similar in *P*– and *P*+ infected cells (Figure 2I), consistent with the former’s ability to lysogenize [8, 11]. Tracking the progression of cell states during infection revealed the transient nature of the “failed lytic” state, as manifested in a decrease in the fraction of lytic-quadrant cells later in infection (Figure S11). This contrasts with the irreversible commitment suggested by *P*+ infection data (Figure 2G). Overall, the analysis of infection by non-replicating phages substantiates the role of active viral replication in enabling an MOI-driven binary choice of cell fate.

### Spatial clustering of mRNA delineates individual phage genomes in the cell

Having characterized the transcription profile of single cells during the infection process, we next sought to zoom-in further and identify the transcriptional activity of the individual phages coinhabiting each cell. Several lines of evidence suggest that single-phage activity may not simply mirror the whole-cell trajectory, but rather exhibit a degree of individuality [9, 10, 30]. However, this potential individuality remained hidden in previous characterization of the post-infection decision.

In earlier work, we used simultaneous detection of a genetic locus and its encoded mRNA to quantify active transcription from endogenous genes in *E. coli*, utilizing the physical proximity of the nascent RNA and its DNA template [17]. Using this approach, we measured the amount of nascent RNA at sister gene copies present during the chromosome replication cycle [17]. Here, we reasoned that a similar approach can be applied to profile the transcription from infecting phages’ genomes, by combining spatial information from mRNA labeling (par-seqFISH) with the two genome labeling schemes used for measuring MOI: EdU-Click, which detects the incoming phage genomes [19], and ParB-*parS* tagging of all viral copies present [11](Figure 1C; see METHODS). And, indeed, as in the case of chromosomal genes [17], lambda transcripts can be partitioned into gene-proximal and distal populations (Figure 3A, Figure S12), enabling the distinction between nascent and mature RNA and associating the former with its encoding phage genome. Unfortunately, we found that the two genome labeling signals degraded during the par-seqFISH procedure (Figure S13). As a result, relying on those signals would only enable us to detect a fraction of intracellular phages, providing inadequate sampling of single-phage behavior. The approach thus had to be amended to allow reliable phage identification.

**Figure 3.**
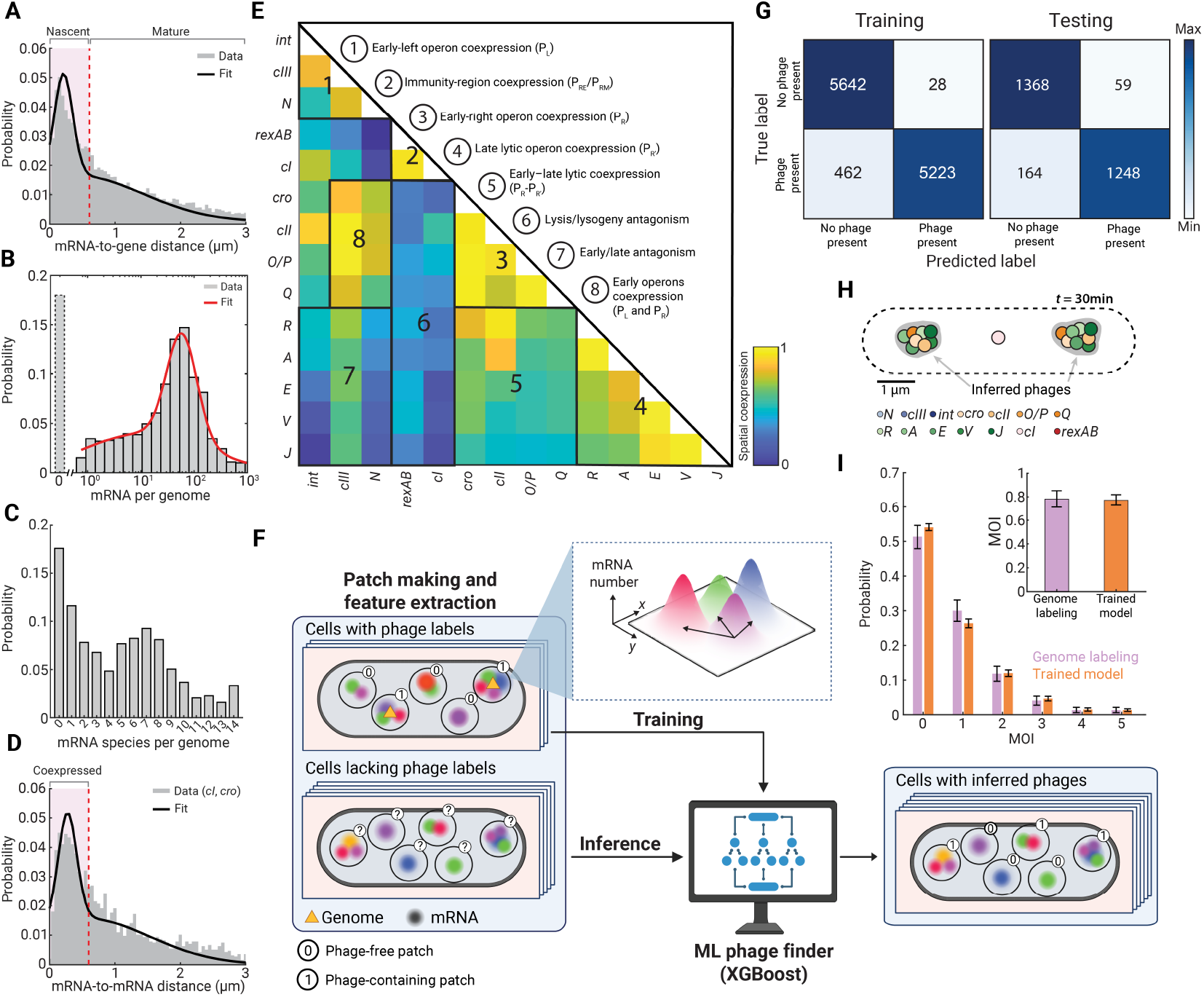
Spatial clustering of mRNA delineates individual phage genomes in the cell. (A) **The distribution of distances between all RNA spots and the nearest labeled phage genome**. Gray, data from *n* = 889 cells (24,132 pairs of mRNA and EdU-Click-labeled genome). Black, fit to a sum of two Gaussians, corresponding to nascent (proximal) and mature (distal) RNA populations. The dashed red line indicates the distance threshold separating the species (*µ* + 2*σ* = 0.61 µm). (B) **The distribution of nascent mRNA per phage**. Gray, data from *n* = 1,441 phage genomes (detected using EdU-Click labeling) in 889 cells; for each phage, the total number of nascent transcripts (of any gene) was tabulated. Red, fit to a sum of two log-normal functions, serving as guide to the eye. (C) **The distribution of mRNA species per phage**. Same dataset as in panel B; for each phage, the number of distinct (i.e., corresponding to different genes) nascent mRNA molecules was tabulated. (D) **The distribution of distances between nearest mRNA molecules of different species**. Gray, the distances between each spot of *cro* mRNA and the nearest *cI* mRNA spot in the same cell was recorded in 520 cells (*n* = 2,557 mRNA-mRNA pairs). See Figure S14 for all gene combinations. Black, fit to a sum of two Gaussians. The dashed red line indicates the distance cutoff (600 nm) used to classify *cI*–*cro* pairs as coexpressed from the same phage and to set the coexpression radius used in panel E. (E) **Spatial coexpression of phage transcripts**. The normalized spatial coexpression between pairs of targeted transcripts was calculated as follows: For each gene pair, we quantified the within-radius neighbor fractions *M*_1_ and *M*_2_, defined as the fractions of spots of each gene whose nearest partner from the other gene lies within 600 nm, and calculated the symmetric coexpression score 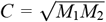. Scores were then normalized row-wise to [0, 1] and symmetrized as (*M* + *M*^T^)/2. For each gene-pair entry in the heat map, *n* = 257 – 1,304 cells (3,128 – 41,703 mRNA pairs). Numbered regions highlight coexpression or antagonistic modules as indicated. (F) **A machine-learning approach to infer the positions of intracellular phage genomes based on RNA spatial features**. Cells were subdivided into overlapping circular patches of radius 300 nm, and each patch was characterized by the mRNA copy numbers, identity, and spatial positions. Patches from cells with genome labels (top left) were used to train an Extreme Gradient Boosting (XGBoost) classifier to distinguish phage-containing from phage-free regions. The trained model was then applied to all cells to infer the positions of phage genomes (right). Created with BioRender.com. (G) **Evaluation of the trained model**. Confusion matrices for training and testing data, indicating strong agreement between predicted and true patch labels derived from RNA content and from genome labeling (training: 11, 355 patches; testing: 2, 839 patches; 889 cells total). (H) **Model-based inference of single-phage transcription profiles**. A representative cell (*t* = 30 min) illustrating application of the trained XGBoost model to infer single-phage transcription from spatial RNA features. Inferred phage locations (gray) are indicated within the segmented cell boundary (dashed). (I) **The trained model recapitulates the MOI distribution obtained from genome labeling**. The single-cell MOI was obtained from EdU-Click genome labeling before par-seqFISH (purple; same dataset as in Figure 1C) and from the trained model, corrected for transcriptionally silent phages (orange; *n* = 485 cells, pooled from 5 and 20 min; see METHODS). Error bars for the genome-labeling data indicate the standard error of the measured fraction *p* in each MOI bin, estimated as 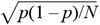; error bars for the model inference are defined as described in METHODS. Inset, the mean MOI values obtained using the two approaches, with error bars denoting mean ± SEM for genome labeling and propagated uncertainty for the model-inferred estimates (see METHODS).

The simultaneous detection of multiple mRNAs in the par-seqFISH procedure presented us with a potential route to delineating the transcription activity of individual phages in the cell, even in the absence of a genome label. To do so, we rely on the proximity of RNAs co-transcribed from the same phage genome: First, applying the approach of [17] to the surviving EdU-Click-labeled genomes, we found that ≈ 82% of phages are transcriptionally active (defined as having mRNA colocalized with the genome), with ≈ 70% of phages transcribing 5 or more mRNA molecules (Figure 3B). Furthermore, ≈ 70% of phages are simultaneously transcribing two or more different genes (Figure 3C). These observations led us to hypothesize that the colocalization of multiple transcripts can be used to identify individual phage genomes and quantify their transcriptional activity. To evaluate the feasibility of this idea, we examined the distribution of distances between mRNA molecules of different genes. Similarly to the mRNA-to-gene distances (Figure 3A), mRNA-to-mRNA distances frequently indicated the presence of two populations, with the proximal one consistent with co-transcription (Figure 3D, Figure S14). The proximal peak was more prevalent in gene pairs expected to be co-transcribed, such as those expressed from the same promoter or during the same developmental stage (Figure 3E). Conversely, mRNAs of genes expressed at different times or under different fate choices were dominated by the distal population, indicating low propensity to be co-transcribed from the same phage (Figure 3E). It is thus plausible that the pairwise distances of different mRNA may be used to identify the positions of (unlabeled) phage genomes and infer their transcriptional profile.

To achieve this goal, we trained an Extreme Gradient Boosting (XGBoost [31]) machine-learning algorithm to infer the positions of phage genomes from the identities, positions, and intensities of mRNA signals in each cell (Figure 3F). The algorithm scans 300 nm-radius “patches” covering each cell and infers, based on the above input properties, whether a phage genome is present or absent from the patch. To train the algorithm, we utilized surviving Click-labeled genomes as “phage-present patch” ground truth. Training on 11,355 curated patches yielded 96% accuracy, and subsequent testing on 2,839 previously unseen patches achieved 92% accuracy for patch-level labels (Figure 3G). Comparing the classification results of the algorithm to the full set of cells in which the genome label was present suggested a high degree of agreement, as indicated by a rand index [32, 33] of 0.99 ± 0.06 (*mean* ± *SD, n* = 703 cells) (Figure S15). Finally, we compared the XGBoost-inferred MOI to the values measured using EdU-Click labeling before par-seqFISH (i.e., before deterioration of the signal, Figure 1C), and found good agreement between the estimates (when accounting for the fraction of silent genome copies) (Figure 3I). Overall, the data thus suggested that the inference algorithm successfully identifies the spatial positions of transcriptionally-active phage genomes.

### Single-phage profiling illuminates viral individuality during infection

We next used the trained algorithm to profile the transcriptional activity (namely, the amount of nascent mRNA for each gene) of individual phages during the infection process. When averaged over all phages in all cells, the observed kinetics closely resembled the averaged whole-cell data (Figure 4A, Figure S16). In addition, applying PCA to the single-phage data yielded principal vectors that resembled those of the single-cell data (Figure 4B, Figure S16). These similarities are unsurprising: While it is true that the phage profiles represent nascent mRNA species, as opposed to the total (nascent plus mature) species captured at the whole-cell level, the short mRNA lifetimes result in comparable levels of the two species in the bacterial cell [17, 24], including for phage genes during lambda infection [11]. However, whereas the phage- and cell-based data is thus consistent, it is the potential divergence between the two observables that is potentially informative, and what we sought to examine next.

**Figure 4.**
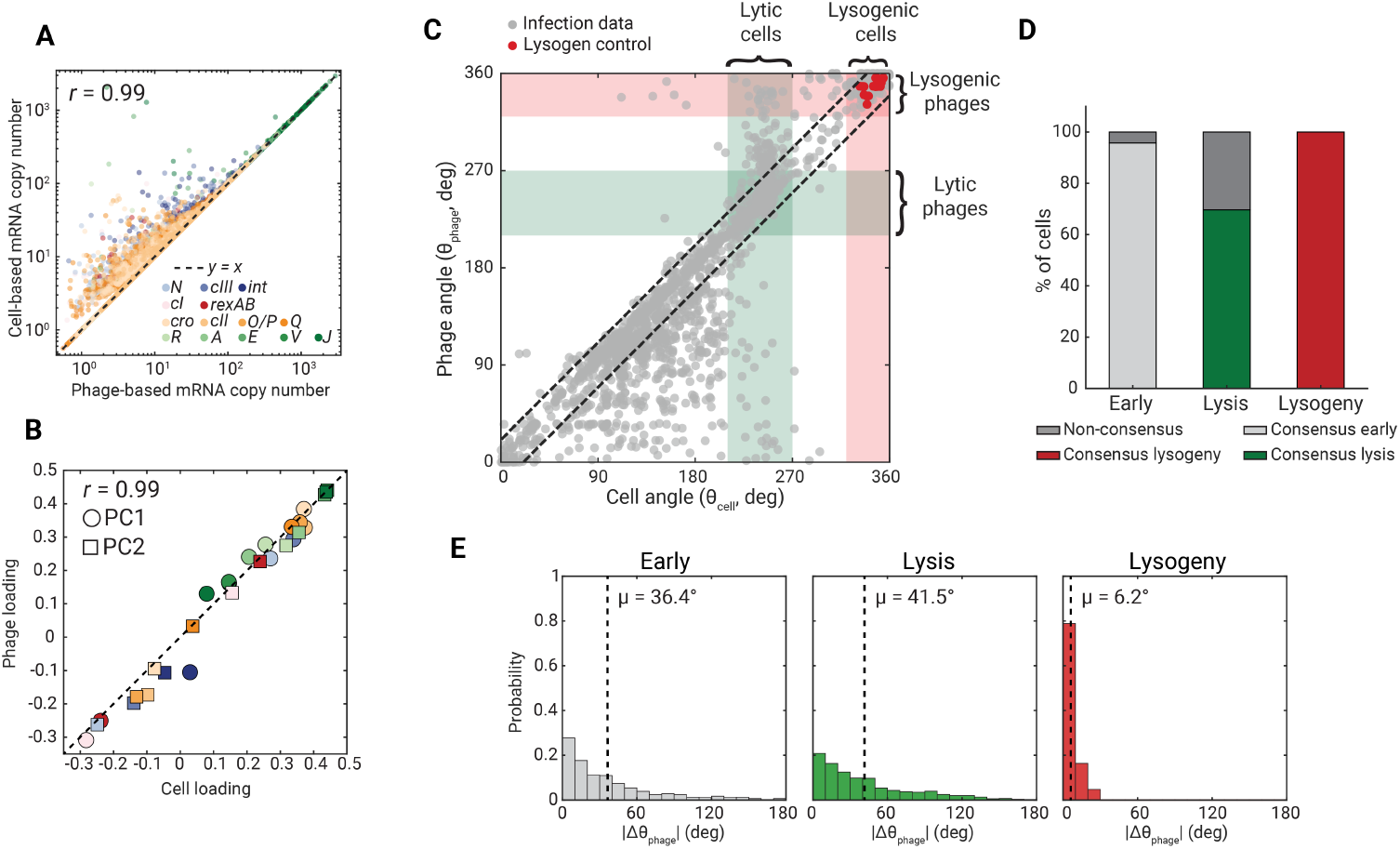
Single-phage profiling illuminates viral individuality during infection. (A) **Comparison of whole-cell and single-phage mRNA copy numbers**. Points, mRNA copy number measurements in individual cells (*n* = 1,463 cells), with *x* values indicating the sum over all inferred phages in the cell and *y* values the corresponding whole-cell estimate. Genes are colored as indicated in the legend. *r*, Pearson correlation coefficient between the two estimates. (B) **Comparison of PCA eigenvectors from cell-based and phage-based analyses**. The loadings of the first two principal component vectors (PC1, circles; PC2, squares) are plotted, with *x* values indicating the cell-based PCA and *y* values the phage-level PCA. Genes are colored as in panel A. Dashed line, *y* = *x. r*, Pearson correlation between cell and phage loadings (C) **Cell angle (***θ*_**cell**_**) versus phage angle (***θ*_**phage**_**) for individual phages**. Each point corresponds to a single phage (*n* = 2, 562). Infection samples are shown in gray and control lysogens in red. Vertical shaded bands indicate the cell-angle windows used to define lytic and lysogenic cells; horizontal bands indicate the analogous phage-angle windows. Dashed lines indicate*y* = *x* ± 20^◦^. (D) **Degree of intracellular consensus at the different developmental stages**. Bars, for cells classified (based on *θ*_cell_) as “early” (*n* = 1,088 cells), “lysis” (*n* = 342 cells), or “lysogeny” (*n* = 309 cells), the percentage of cells in which all detected phages share the corresponding phage-angle fate or exhibit mixed phage classifications (non-consensus). (E) **The distributions of pairwise differences in** *θ*_**phage**_ **within cells**. For each cell class (early, gray; lysis, green; lysogeny, red), the probability distribution of the absolute difference in phage angle, |Δ*θ*_phage_|, between all phage pairs in the same cell (early, *n* = 882 phage pairs; lysis, *n* = 458 phage pairs; lysogeny, *n* = 379 phage pairs). Dashed lines mark the mean |Δ*θ*_phage_| for each class.

In earlier work, we used live-cell microscopy to measure the frequency of lysogenization as a function of the MOI (quantified using fluorescently labeled phage capsids) and the size of the infected cell [9, 34]. Our original expectation was that this frequency (*f*_lys_) would scale with the ratio of MOI to cell volume, 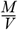. This ratio approximates the concentration of viral gene products in the cell, which, in turn, is sensed by the lambda-encoded decision circuit to determine the developmental trajectory [35]. We were surprised to find that, rather than scaling directly with 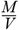, the probability of lysogeny exhibited an added exponential (power) dependence on the MOI, i.e., 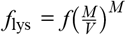. To explain this relation, we invoked an added requirement for cell fate determination: To achieve lysogeny, all phages present in the infected cell must choose that fate; in the absence of consensus, the lytic route is followed. This simple scenario successfully captured the observed phenomenology [9]. However, it remained speculative absent the ability to characterize the expression profiles of individual phages within the cell and identify their developmental preference.

Our new ability to profile the transcription of individual coinfecting phages allowed us to address the degree of viral consensus during the lysis/lysogeny decision. To do so, we plot for each cell the value of *θ*_cell_ (indicating choice of fate, Figure 2D above) versus the values of “phage angles” (*θ*_phage_), defined identically to *θ*_cell_ but using individual phages’ expression profile (hence indicating whether each phage is expressing lytic or lysogenic genes), for all phages present in that call (Figure 4C, Figure S17). In the absence of phage individuality, we expect the data to lie approximately on the diagonal, representing a consensus among all phages within each infected cell. And, indeed, a near-diagonal population is clearly observed, with ≈ 76% of phages found within ±20^◦^ of the diagonal. However, a significant fraction of the population resides outside this “intracellular consensus” region, and the deviation from consensus varies with the developmental state of the cell: In “early infection” cells (*θ*_cell_ = 0^◦^ – 210^◦^), phages are often found off the diagonal (≈ 28% of phages >20^◦^ away from it), likely indicating lack of synchronization in the developmental progression of individual phages prior to fate commitment. Once cells leave this early stage and converge towards one of the two eventual outcomes, a striking difference is observed: In lytic cells, phages are often found far from the diagonal (≈ 24% of phages >20^◦^ away from it), with the majority of those (≈ 15%) on the lysogenic side (higher value of *θ*_phage_) of the plane. Lysogenic cells, in contrast, appear much more consensual (≈ 4% at > 20^◦^ away from the diagonal). Specifically, no lytic-profile phages (0%) are found in cells pursuing lysogeny (similar to the value in the lysogenic control, 0%). The degree of intracellular consensus at the different developmental stages is summarized in Figure 4D. A similar trend is found when examining the distributions of angular differences between pairs of phages within the cell, Δ*θ*_phage_ (Figure 4E), which again indicate near consensus inside lysogenic cells, but lack of such consensus in lytic ones. Taken together, the profiling of single-phage transcription is thus consistent with the earlier inference of [9], namely, that the choice of lysogeny requires a consensus among coinfecting phages, whereas any deviation from unanimity results in lysis.

## DISCUSSION

Despite the paradigmatic role played by the lambda decision [4, 20] and its being one of the best characterized living systems [4, 20, 36, 37], an empirical characterization of the diverging transcriptional trajectories following infection has proven elusive. While such divergence is clearly tractable in the simpler two-gene maintenance “switch” [38–40], whose state can be easily toggled using, e.g., a temperature-sensitive repressor [38–40], true infection, during which the full viral circuit comes into play, presents a considerably more challenging case [23, 41]. Here, to characterize single-cell trajectories, we employed an infection protocol in which the steps of phage adsorption, entry, and intracellular growth are optimized to temporally synchronize the infected cells, and to achieve a distribution of single-cell MOI that ensures lysis and lysogeny are represented in comparable numbers [9, 11]. Applying par-seqFISH to time-lapsed samples following infection reveals the single-cell signatures of the two developmental trajectories. PCA then allows mapping the data into two interpretable dimensions, corresponding to the progression of viral genome transcription and the divergence between lysis and lysogeny. The two eventual cell fates map to different quadrants of this two-dimensional plane (Figure 2C).

Whereas this cell-level analysis coheres with the canonical view of lambda infection, our ability to profile the transcriptional activity of individual phages within each cell reveals behavior that remains unexplained by the current narrative. Specifically, we find that lytic cells are often inhabited by phages whose transcription profile indicates the choice of lysogeny. In lysogenic cells, in contrast, a consensus choice among all phages is found (Figure 4E). Crucially, the data does not allow us to identify the direction of causality, namely, whether cell lysogenization requires phage unanimity, leads to it, or perhaps the two features are not causally related. Nevertheless, interpreted in light of the earlier inference of [9] and [10], the observed phenomenology lends credence to the idea that phages maintain a degree of individuality during cell fate determination, and that a unanimous decision among all coinfecting phages is required for the establishment of lysogeny.

What would allow coinfecting phages to make individual decisions? While the bacterial cytoplasm was traditionally considered well mixed [42], evidence to the contrary has been steadily accumulating [10, 43–45]. It is thus tempting to speculate that the genetic activity of individual phages coinhabiting the cell can be decoupled, at least partly. Such decoupling may come about in multiple ways, for example, through cis-preferential activity of phage transcription factors, a feature that has been predicted theoretically [46] and reported in early studies of lambda and other phages [47–49].

Beyond its particulars, the infection scenario studied here can begin to illuminate the broader case: The genetic elements that drive cell function exist, almost invariably, in multiple distinct copies in each cell. The number of copies changes over time, e.g., as part of the cell replicative cycle [50], and their activity may be correlated—positively or negatively—in ways that are often poorly understood [17, 51]. The corresponding copy numbers and their temporal kinetics are accentuated further under abnormal scenarios such as genomic copy number variations or viral infection [52]. Giving explicit consideration to these discrete genetic copies is critical to our ability to understand, and predict, cellular behavior. As in other instances, lambda infection may serve as our starting point for the elucidation of new biology.

## METHODS

### Chemical reagents, growth media, and buffers

Chemical reagents, growth media, and buffers used in this study are listed in Table S2 and Table S3. Supplements to growth media are described for each experiment, as appropriate. Agar plates and soft agar were prepared by adding 1.5% and 0.7% (w/v) agar, respectively [53]. All media were sterilized by autoclaving at 121^◦^C for at least 25 min using a liquid cycle.

### Probe design

#### Primary probes

Primary probes were designed following [15]. Each probe contained a 30-nucleotide sequence complementary to the target mRNA, flanked by two 15-nucleotide secondary hybridization sites at both the 5^*′*^ and 3^*′*^ ends (the design of secondary probes is described in Readout probes). The mRNA-complementary regions of primary probes were designed using the ProbeDealer software [54], with design parameters as described in [15]. Briefly, probes were selected based on a GC content of 45–65%. Sequences containing more than four consecutive identical nucleotides were excluded. To minimize non-specific binding, candidate probes were aligned against the genomes of *Escherichia coli* strain MG1655 (NCBI accession: U00096.3) and bacteriophage lambda (NCBI accession: NC_001416) using BLAST (NCBI BLAST, 2025) [55], and any probe with potential off-target hybridization of at least 18 contiguous nucleotides was discarded. For each target gene, 6 to 20 non-overlapping probes were selected, depending on gene length. In cases where target genes were too short to accommodate a sufficient number of probes, adjacent genes transcribed from the same promoter were grouped and targeted collectively, specifically: {*cIII, kil, gam*}, {*rexA, rexB*}, and {*O, P*}.

In total, 178 primary probes were designed, targeting transcripts from 14 distinct regions of the lambda genome and encompassing 18 genes: *int, gam, kil, cIII, N, rexB, rexA, cI, cro, cII, O, P, Q, R, A, E, V*, and *J* (Figure 1D). Gene selection was performed to ensure adequate representation of the lambda genome in terms of transcription units and functional gene categories [20]. The full list of primary probe sequences is provided in (Table S4). All primary probe sequences were ordered as an oligonucleotide pool from Integrated DNA Technologies (IDT) at a scale of 50 pmol per oligo.

#### Ribo-tag probes

Probes targeting the 16S ribosomal RNA (Ribo-tag probes) were designed using the approach of [15]. Ribo-tag probes were directed against a conserved region of the *E. coli* 16S rRNA, and adhered to the same selection criteria applied to the primary probes, with an rRNA-complementary region length of 28 nucleotides. Each Ribo-tag probe was flanked by one secondary hybridization sequence at the 5^*′*^ end and one at the 3^*′*^ end, selected from a set of six sequences (described in Readout probes) to enable multiplexed detection. All Ribo-tag probes are listed in Table S5.

#### Readout probes

Readout probes, each 15 nucleotides in length, were adopted from a subset of the readout probe library reported in [15]. These sequences were generated by random permutation of the four nucleotides (A, T, G, and C), followed by retention of sequences with GC content between 40% and 60% and exclusion of homopolymer stretches [15]. From this pre-validated library, we selected a subset of readout probes and further validated them through genome-wide off-target screening and cross-hybridization filtering, as described for primary probes (see Primary probes). The reverse-complement sequences of the validated readout probes were appended to the 5^*′*^ and 3^*′*^ ends of the primary probes (see Primary probes and Ribo-tag probes), with each gene assigned a specific readout probe. The complete set of assignments is provided in Table S4 and Table S5. Readout probes, each labeled at the 5^*′*^ end with Alexa Fluor 647, TAMRA, or Alexa Fluor 594, were ordered as high-performance liquid chromatography (HPLC)-purified oligonucleotides from IDT.

### Bacterial strains, phages, and plasmids

All strains used in this study are listed in Table S1. *E. coli* strains MG1655 and LE392 were used, along with two bacteriophage lambda strains: *λ*_TY5_ (*λ cI857 Sam7 stf* ::P1*parS-kan*^R^) and *λ*_TY11_ (*λ cI857 Pam80 stf* ::P1*parS-kan*^R^) [11].

*λ*_TY5_ carries an amber mutation in the *S* gene, which encodes the holin protein required for cell lysis. This mutation prevents lysis in the wild-type host MG1655 [37]. *λ*_TY11_ carries an amber mutation in the *P* gene, which is essential for phage DNA replication; this mutation blocks replication in wild-type host MG1655 [37]. In addition to these mutations, both *λ*_TY5_ and *λ*_TY11_ carry a P1-derived *parS* sequence inserted at the *stf* locus [11].

To label phage genomes in infected cells, we used the ParB–*parS* system. A fluorescently tagged ParB protein (CFP–ParB), expressed from plasmid pALA3047 (a gift from Stuart Austin), binds specifically to the *parS* site, allowing visualization of phage DNA [11]. In addition, *λ*_TY5_ was labeled metabolically using 5-ethynyl-2′-deoxyuridine (EdU), followed by detection through click chemistry, as described in Production of EdU-labeled *λ*_TY5_ and Click labeling and imaging.

### Phage preparation

#### Production of phage *λ*_**TY11**_

Crude phage lysates were prepared using a protocol adapted from [56, 57]. Briefly, an overnight culture of LE392 *λ*_TY11_ lysogens was diluted 1:1000 into LBGM medium (Table S2) in a baffled Erlenmeyer flask and incubated at 30^◦^C with shaking at 180 rpm. Upon reaching OD_600_ ≈ 0.4, the culture was transferred to a 42^◦^C water bath and shaken at 180 rpm for 15 min, followed by incubation at 37^◦^C with continued shaking for approximately 1 h until the OD_600_ dropped below 0.05.

After lysis, 5% (vol/vol) chloroform was added to the culture, and the sample was incubated at room temperature for 15 min. The lysate was then centrifuged at 4000×g for 10 min at 4^◦^C to pellet cell debris. The clear supernatant was collected, supplemented with 0.3% (vol/vol) chloroform, and stored at 4^◦^C until use. Phage titers were determined by standard plaque assays on NZYM agar plates (Table S2) using LE392 reporter cells, yielding titers of 10^10^–10^11^ plaque-forming units per mL (PFU/mL).

#### Production of EdU-labeled *λ*_TY5_

EdU-labeled *λ*_TY5_ phages were produced by infecting MG1655 host cells in the presence of EdU, using a protocol adapted from [19]. Overnight cultures of MG1655 were diluted 1:1000 into 25 ml of LBGM (Table S2) in a baffled Erlenmeyer flask and incubated at 37^◦^C with shaking at 220 rpm. Upon reaching OD_600_ ≈ 0.4, cells were harvested and concentrated 10-fold by resuspending the pellet in 1 mL of LBMM (Table S2). For infection, 200 µL of concentrated cells (approximately 1×10^9^ cells/mL) were mixed with 20 µL of *λ*_TY5_ phage stock (2×10^10^ PFU/mL) to achieve an MOI of 2. After 20 min of adsorption at 37^◦^C, the 220 µL infection mixture was diluted into 25 mL of fresh LBGM (prewarmed to 37^◦^C) containing 100 µM EdU and incubated at 37^◦^C with shaking at 180 rpm for 3 hours. Following incubation, cells were harvested by centrifugation, washed twice with 1×PBS to remove unadsorbed phages, and resuspended in 500 µL of LBMM. Lysis was induced by adding chloroform to 5% final concentration and incubating at room temperature for 20 min with periodic vortexing. Cell debris was removed by centrifugation (4500×g, 5 min), and the lysate was stored with 0.3% (vol/vol) chloroform at 4^◦^C. Phage titers were determined by plaque assays on NZYM agar plates (Table S2) using LE392 reporter cells, typically yielding titers of 10^10^–10^11^ PFU/mL.

### Phage infection and sample preparation

#### Phage infection

The infection protocol was adapted from [11], with adjustments to culture volumes to accommodate downstream par-seqFISH processing. An overnight culture of *E. coli* MG1655 carrying plasmid pALA3047 was diluted 1:1000 into 90 mL of LBMM supplemented with 10 µM IPTG and grown at 30^◦^C with shaking at 220 rpm in a 1 L baffled Erlenmeyer flask. In parallel, MG1655 *λ*_TY9_ lysogens carrying plasmid pALA3047 (serving as the lysogen control sample) was diluted 1:1000 into 15 mL of LBMM supplemented with 10 µM IPTG and grown under identical conditions. Upon reaching OD_600_ ≈ 0.4, 10 mL from each culture were retained as uninfected and lysogenic control samples, and immediately processed according to the fixation procedure described in Fixation and permeabilization. From the non-lysogenic culture, 60 mL was collected for infection, divided equally into two 50 mL centrifuge tubes (Corning) and centrifuged at 3,000 rpm for 10 min at 4^◦^C. The supernatant was decanted, and each cell pellet was resuspended in 300 µL of fresh, ice-cold LBMM supplemented with 10 µM IPTG, resulting in a ∼100× concentration relative to the overday culture (approximately 6×10^9^ cells/mL). The resuspended pellets were then pooled.

To determine viable cell concentration in the 100× suspension, 10 µL of concentrated host cells was diluted 10^–7^ in 1×PBS, and 100 µL was plated on LB agar plates in three replicates. Colony-forming units (CFU) were counted after incubation at 30^◦^C overnight, and the concentration (CFU/mL) of the original suspension was calculated.

To begin infection, 550 µL of concentrated host cell suspension was mixed with a volume of phage stock (prepared as described in Phage preparation) adjusted to reach an MOI ≈ 1 (typically, < 100 µL). The infection mixture was incubated on ice for 30 min to allow adsorption, followed by incubation at 35^◦^C for 5 min to trigger phage DNA injection. Subsequently, 600 µL of the infection mixture was diluted 1:1000 into 600 mL LBGM (prewarmed to 30^◦^C) supplemented with 10 µM IPTG, divided into two 2 L baffled Erlenmeyer flasks, and incubated at 30^◦^C with shaking at 180 rpm (the beginning of incubation is defined as *t* = 0 min). At specified time points (0, 1, 2.5, 5, 7.5, 10, 15, 20, 30, 45, and 60 min), 40 mL aliquots were collected and processed as described in Fixation and permeabilization.

#### Lysogenization assay

To measure the fraction of lysogens in the infected population, 1 mL samples were collected from each infection flask at *t* = 45 min after dilution into LBGM (see Phage infection). Each sample was diluted 10^–4^ in 1×PBS, and 100 µL of the diluted suspension was plated on LB agar plates, supplemented with 50 µg/mL kanamycin, in two replicates per flask. Plates were incubated at 30^◦^C overnight, and colonies were counted. The lysogenization frequency in the culture was calculated as the ratio of kanamycin-resistant CFU to the total CFU measured in Phage infection. Uncertainty was estimated by nonparametric bootstrapping [58, 59] across biological replicates (1,000 resamples with replacement); the lysogenization frequency was recalculated for each resample, and error bars report the bootstrap standard error (standard deviation of the bootstrap distribution).

#### Fixation and permeabilization

Fixation and permeabilization procedures were adapted from [11]. For infected samples at each time point, 40 mL culture was mixed directly with 10 mL of 18.5% formaldehyde in 5×PBS. For uninfected and lysogen controls, 10 mL of culture was mixed with 2.5 mL of the same fixative solution. In all cases, this resulted in a final concentration of 3.7% formaldehyde in 1×PBS. Samples were incubated on a nutator at room temperature for 30 min. Following fixation, cells were washed twice with 1 mL 1×PBS by centrifugation at 600×g for 3.5 min. For samples collected at *t* = 5 and 20 min, fixed cell pellets were resuspended in 1 mL of DEPC-treated water, and a 250 µL aliquot was reserved for single-cell MOI determination (performed as described in MOI determination by microscopy), and the remaining 750 µL was used for permeabilization. For all other time points, cell pellets were resuspended directly in 750 µL of DEPC-treated water. For permeabilization, 750 µL of cell suspension was mixed with 250 µL of 100% ethanol and incubated on a nutator at room temperature for 1 h.

#### MOI determination by microscopy

To quantify the number of intracellular phage genomes per cell (single-cell MOI), fixed samples collected at *t* = 5 and 20 min (see Fixation and permeabilization) were processed as follows. From each 250 µL sample, 50 µL was pelleted and resuspended in 10 µL 1×PBS, then mounted on coverslips under 1% agarose pads for imaging to visualize ParB-*parS*-labeled phage genomes, as described in [11]. For infections with EdU-labeled phages (see Production of EdU-labeled *λ*_TY5_), the remaining 200 µL from each 250 µL underwent click labeling using the Click-iT EdU Cell Proliferation Kit (Fisher Scientific) according to the manufacturer’s protocol before imaging. Imaging was performed as described in Click labeling and imaging. Diffraction-limited fluorescent foci were counted in individual cells, and the number of foci per cell was interpreted as the single-cell MOI (Figure 1C, Figure S1).

### Parallel sequential fluorescence in situ hybridization (par-seqFISH)

#### Coverslip functionalization

Glass-bottom flow chamber (ibidi) was used for sample mounting. Prior to use, the glass surface was coated with 0.01% poly-L-lysine solution (Sigma-Aldrich) for ≥1 h at room temperature, following the manufacturer’s protocol [60]. Following incubation, the chambers were rinsed with 1 mL DEPC-treated water and allowed to air-dry completely.

#### Primary probe hybridization

The par-seqFISH protocol was adapted from [15, 16]. Approximately 10^8^ permeabilized cells from each sample were collected by centrifugation at 2000 rpm for 7 min and resuspended in 10 µL nuclease-free water. 10 µL of a sample-specific Ribo-tag probes (50 nM stock concentration; final concentration: 10 nM), each consisting of a pair of primary probes carrying distinct secondary sequences (Table S5), was added to the cell suspension, followed by the addition of 30 µL of prewarmed primary hybridization buffer (50% (vol/vol) formamide, 10% (wt/vol) dextran sulfate, 2×SSC). Samples were incubated at 37^◦^C for >16 h to allow probe hybridization. Following hybridization, samples were washed twice with 100 µL of wash buffer (55% (vol/vol) formamide, 0.1% (vol/vol) Triton X-100 in 2×SSC) by centrifugation at 8000 rpm for 5 min. A subsequent 30 min incubation at 37^◦^C in 100 µL of wash buffer was performed to remove nonspecifically bound probes. Samples were then washed twice with 100 µL of 2×SSC to remove residual formamide, and pooled in equal volumes into a single Eppendorf tube. The pooled cells were pelleted and resuspended in 100 µL of nuclease-free water. For hybridization with primary gene probes, 10 µL of the pooled cell suspension was mixed with 10 µL of the gene-probe library (50 nM stock concentration per probe; 10 nM final concentration per probe) and combined with 30 µL of prewarmed primary hybridization buffer. Hybridization was performed at 37^◦^C for >16 h. Post-hybridization washes followed the same protocol used for Ribo-tag probe hybridization. The final sample was resuspended in 200 µL 1×PBS, and 40 µL of the cell suspension was gently introduced into a poly-L-lysine–coated flow chamber (prepared as described in Coverslip functionalization). Cells were allowed to sediment and adhere to the surface during a 15 min incubation at room temperature. Unbound cells were removed by washing the chamber with 1 mL of nuclease-free water.

#### Microscopy setup

Imaging was performed using an inverted fluorescence microscope (Eclipse Ti2-E, Nikon) equipped with a motorized stage (TI2-S-SE-E, Nikon), a universal specimen holder, an LED illumination source (X-Cite XYLIS), and a CMOS camera (Prime 95B, Photometrics). A 100× oil-immersion phase-contrast objective (CFI60 Plan Apo *λ*, NA 1.45, Nikon) was used for all acquisitions. Fluorescence detection was performed using the following filter sets: CFP (Nikon, 96341), Narrow Cy3 (Chroma, SP102v1), Narrow Cy5 (Chroma, 49307), and a custom Alexa Fluor 594 set (Omega Optical) comprising an excitation filter (590±10 nm), a 610 nm long-pass dichroic beamsplitter, and an emission filter (630±30 nm). Image stacks were acquired at five z positions (focal planes) with steps of 300 nm. Imaging was performed across multiple fields of view per sample (typically 25–36), to image a total of 2000–4000 cells.

#### Sequential secondary probe hybridization and imaging cycles

The flow chamber with immobilized cells (prepared as described in Primary probe hybridization) was connected to a fluidics system (Elveflow, seqFISH pack) for buffer exchange and probe delivery. Regions of interest were manually identified using phase-contrast imaging. The initial imaging round included the acquisition of phase-contrast images (100 ms exposure) for cell segmentation, followed by imaging in the CFP channel (200 ms exposure) to detect CFP–ParB foci. Subsequent imaging rounds involved iterative cycles of readout probe hybridization, washing, and fluorescence imaging, adapted from [15, 16]. Each hybridization round involved three color-distinct secondary readout probes labeled with TAMRA, Alexa Fluor 647, or Alexa Fluor 594. Probes were diluted into ethylene carbonate (EC) hybridization buffer (10% (vol/vol) ethylene carbonate, 10% (wt/vol) dextran sulfate, 4×SSC) and introduced into the flow chamber at a rate of 50 µL/min, followed by 20 min incubation. Post-hybridization washes were performed by flowing ∼1 mL of wash buffer (10% (vol/vol) formamide, 0.1% (vol/vol) Triton X-100 in 2×SSC) through the chamber at 200 µL/min to remove unbound probes, followed by a rinse with ∼1 mL of 4×SSC at the same flow rate. Phase-contrast images were acquired first (100 ms exposure), followed by fluorescence imaging in the Narrow Cy3 (500 ms; TAMRA), Narrow Cy5 (500 ms; Alexa Fluor 647), and Alexa 594 (500 ms; Alexa Fluor 594) channels. Following each imaging round, readout probes were stripped by flowing ∼1 mL 55% wash buffer (55% (vol/vol) formamide, 0.1% (vol/vol) Triton X-100 in 2×SSC) through the chamber at 200 µL/min, followed by a 15 min incubation. The sample was subsequently rinsed with ∼1 mL of 4×SSC at 200 µL/min to remove residual formamide. This process of hybridization, imaging, and probe stripping was repeated for seven sequential rounds, enabling multiplexed detection of 14 target mRNAs across 13 conditions (11 time points, plus lysogen and uninfected controls).

#### Click labeling and imaging

For samples infected with EdU-labeled *λ*_TY5_ phages, following the final round of secondary probe hybridization and stripping, a click reaction was performed to detect the DNA of entrant phage genomes. The Click-iT EdU Cell Proliferation Kit for Imaging, Alexa Fluor 647 dye (Fisher Scientific), was used with modifications to the manufacturer’s protocol to accommodate the fluidics system. Briefly, the click reaction cocktail was freshly prepared and introduced into the flow chamber at a rate of 50 µL/min. Samples were incubated with the reaction mixture for 30 min. Following incubation, the samples were washed with 2 ml of 1×PBS at 200 µL/min to remove unreacted reagents. Phase-contrast images were acquired first (100 ms exposure), followed by fluorescence imaging in the Narrow Cy5 channel (500 ms, Alexa Fluor 647).

### Image analysis

#### Cell segmentation and tracking

Each field of view contained a sequence of images corresponding to the imaging rounds described in Sequential secondary probe hybridization and imaging cycles, and, when applicable, the additional click-labeling imaging described in Click labeling and imaging. In each field of view and for every imaging round, the focal plane was defined as the one with the highest pixel intensity variance [11]. Cell segmentation was performed on these in-focus slices using a U-Net convolutional neural network [61], previously trained on phase-contrast images of *E. coli* acquired under similar imaging conditions [27]. The resulting segmentation masks and the original images were then used as input to Delta 2.0 [62] to track individual cells across sequential imaging rounds within each field of view.

#### par-seqFISH demultiplexing

Cells harvested at each time point were uniquely labeled using specific pairs of Ribo-tag probes (see Primary probe hybridization). For demultiplexing these samples, maximal intensity projection images were generated from the z-stacks for each 16S rRNA readout channel, and background fluorescence was estimated by averaging the intracellular pixel intensity across all segmented cells. This background was subtracted to compute the total fluorescence per cell for each rRNA probe. For classification, the background-corrected intensities of the two 16S probes assigned to each barcode (Table S5) were plotted against one another on a log scale, and cells with high signal in both channels were identified by gating and assigned to the corresponding time point according to the codebook (see Primary probe hybridization and Table S5). Cells that matched multiple time points or none were labeled as “unknown.” Using these criteria, approximately 90% of cells were assigned unambiguously. The false-positive decoding rate was estimated by counting cells assigned to barcode combinations excluded from the experiment and was found to be approximately 0.1%.

#### Spot recognition

Spot recognition was performed using Spätzcells [22] and applied to par-seqFISH-labeled mRNA, ParB-*parS*-labeled phage loci, and EdU-Click-labeled phage genomes. Briefly, the software first identified candidate spots by locating, in each focal plane, two-dimensional local maxima of fluorescence intensity above a user-defined detection threshold, and retained only those that were present in at least two adjacent focal planes (z positions). For each retained spot, the fluorescence profile was subsequently fitted to a two-dimensional elliptical Gaussian. The fitting provided the properties of each spot, including position, spot area, peak height (amplitude of the fitted Gaussian), and spot intensity (integrated volume under the fitted Gaussian). Finally, spots were assigned to individual cells based on the segmentation masks (see Cell segmentation and tracking).

#### Discarding false-positive spots

To remove false-positive spots, such as those resulting from nonspecific binding of probes, we performed a gating procedure following the approach of [11]. A low detection threshold was used during spot recognition (see Spot recognition), ensuring that genuine spots were captured, but also increasing the number of false positives. To filter out these false positives, we compared the 2D scatter plots of peak height versus spot area in the experimental samples to those from a negative control consisting of uninfected host cells. A polygon was manually defined on each 2D plot such that most spots appearing in the negative control were excluded. For mRNA, this gating procedure was performed separately for each gene. For ParB-*parS*-labeled phage loci and EdU-Click-labeled phage genomes, gating was similarly based on their respective spot distributions relative to the negative control. Spots falling outside the gated region were discarded. The accuracy of this gating was confirmed by manual inspection of a subset of images.

#### mRNA quantification

The fluorescence intensity corresponding to a single mRNA molecule was estimated following [11]. Briefly, after removing false-positive spots, we fitted the histogram of spot intensities to a sum of three Gaussians, corresponding to one, two, or three mRNA molecules per spot. The mean of the first Gaussian was taken as the estimated intensity of a single mRNA molecule. Each spot’s measured intensity was then divided by this value to estimate the number of mRNA molecules per spot.

#### Phage DNA localization

For ParB-*parS* and EdU-Click spots, only positional information was used; intensity values were not analyzed. After spot detection and assignment to cells (see Inferring single-phage transcription profile in the presence of genome label), the positions of phage spots were used to infer single-phage transcription profiles by linking individual phage genomes to nearby mRNA spots (see Spot recognition), and as ground-truth training labels for the algorithm that detects phage positions based on mRNA spatial features (see Inferring single-phage transcription profile in the absence of genome label).

### Comparison between par-seqFISH and bulk RNA-seq measurements

To validate mRNA abundance measurements obtained by par-seqFISH, we compared them to bulk RNAseq profiles from synchronized infections performed with the same host strain and the same growth and infection protocol used for par-seqFISH, with adjustments to culture volumes as described in [25]. Bulk RNA-seq datasets are available in GEO under accession numbers GSE298101 (*P*+; [25]) and GSE308589 (*P*–; this work).

To compare expression profiles obtained from par-seqFISH and bulk RNA-seq, we computed the Pearson correlation coefficient between log_10_-transformed expression values (with a small offset, *ϵ* = 0.01, added prior to transformation) across all shared genes and shared time points (Figure S5, Figure S10). This analysis quantified agreement in relative mRNA abundance between the two methods across the infection time course.

### Single-cell analysis

#### Principal components analysis

Principal component analysis (PCA) [26] was applied to par-seqFISH gene-expression data from 14 *λ* genes across all samples, to identify the major axes of variation in the expression pattern. Prior to PCA, cells with zero expression across all measured genes (classified as uninfected) were excluded. For each remaining cell, the total copy number of mRNA per cell was normalized to a fixed target of 10,000 copies per cell, a pseudocount of 1 was added, and the expression values were log normalized (with the natural base) and z-scored [45]. Then, PCA (MATLAB function pca) was performed on the resulting scaled gene-expression matrix, and the first two principal components, which together captured approximately 60% of the variance (Figure S7), were used in downstream analysis.

#### Derivation of cell angle

To quantify the developmental trajectory of individual cells, we computed an angular coordinate from the first two principal components, following the approach of [27–29]. Each cell’s angle was calculated from PC1 and PC2 using the four-quadrant inverse tangent (MATLAB function atan2) and defined as *θ* = atan2(*PC*2, *PC*1), yielding values in –180^◦^–180^◦^ with 0^◦^ aligned to the positive PC1 axis. To align this coordinate with the temporal progression of infection, we rotated the angles so that *θ* = 0^◦^ coincides with the line through the origin that places all lysogenic cells above it while minimizing the number of early-time-point cells above it. The polar angle in the rotated system was used as the cell angle.

### Single-phage analysis

#### Inferring single-phage transcription profile in the presence of genome label

In cells with detected phage genomes (using EdU-Click labeling for *λ*_TY5_ infection or ParB-*parS* labeling for *λ*_TY11_ infection; see Sequential secondary probe hybridization and imaging cycles, Click labeling and imaging, and Phage DNA localization), we followed the analysis procedure described in [17] for each mRNA species. Briefly, we computed the distance from the center of each mRNA spot to the center of its nearest phage genome position within the same cell. Histograms of mRNA-to-genome distances revealed two distinct populations: one localized near the genome and one located further away (Figure S12). To parameterize the distribution, we fitted the distance histograms to a sum of two Gaussians. mRNAs falling within the first Gaussian, centered near the genome, were classified as nascent, while those in the second, more distal Gaussian were classified as mature. Based on this classification, we inferred the transcriptional profile of individual phage genomes by associating each genome with its surrounding nascent mRNAs.

#### Inferring single-phage transcription profile in the absence of genome label

To address the limitations in phage detection due to deterioration of genome-labeling signals during sample preparation (Figure S13), we implemented an Extreme Gradient Boosting (XGBoost) machine-learning algorithm [31] to infer phage locations in the absence of detectable genome label. The model was trained on cells in which a phage genome was detected (see Click labeling and imaging, Phage DNA localization, and Inferring single-phage transcription profile in the presence of genome label) to learn the spatial patterns of nascent mRNAs around these genomes, and then applied to predict phage locations based on mRNA spatial features. Within each cell mask (see Cell segmentation and tracking), we placed circular patches (radius 300 nm) on a grid with 350 nm center-to-center spacing. The 300 nm patch radius corresponds to the average gene-proximal distance observed across all measured mRNAs (Figure S12). As an independent validation of the chosen patch size, when we computed pairwise distances between nearest spots from two distinct mRNA species, we observed a population of molecules colocalized within ∼600 nm, i.e., the chosen patch diameter (Figure S14).

For each patch, we extracted the mRNA copy number of individual spots as well as the set of pairwise distances between spots from different mRNA species. For gene pairs in which at least one species had zero copy number within a patch, we treated the pairwise distance as undefined and set it to 5 µm. For patches with multiple mRNA spots corresponding to the same gene species, we calculated their copy number-weighted center of mass and placed a virtual spot at this position, with total copy number equal to the sum of the original spots.

For cells where phage genome label was detected, we generated patches and initially assigned each a binary label: “1” indicating the presence of a phage genome and “0” indicating its absence. Data preprocessing involved two steps. First, when overlapping patches captured the same set of mRNA spots within a cell, we retained a single representative patch and removed the redundant copies. Second, we examined the distribution of total mRNA counts per patch (Figure S15). The distribution was well described by a sum of two exponential functions, indicating the presence of two regimes: a low-count regime (≤10 transcripts per patch) and a higher-count regime (>10 transcripts per patch). We interpreted the low-count regime as likely arising from mature mRNAs that were spatially proximate by chance, whereas the higher-count regime was consistent with nascent mRNAs clustered near active transcription sites. To incorporate this distinction into the training data, patches initially labeled 0 (no genome detected) but with total counts >10 were reassigned to label 1, acknowledging that high local mRNA concentrations indicate probable phage presence.

To ensure robust model training, we balanced the dataset to achieve equal representation of patches with label 0 (phage-absent) and label 1 (phage-present), then partitioned the data into training (80%) and testing (20%) sets. The XGBoost classifier achieved 95% accuracy on the training dataset and maintained 90% accuracy on the test set (Figure 3G). The close agreement between training and test accuracies suggests that the model generalized well without substantial overfitting [63].

The trained model was applied to all cells to infer phage locations and reconstruct phage transcription profiles. After prediction, overlapping patches that shared any mRNA spot were merged into one phage entity if at least one of those patches was predicted to be genome-present. To validate these inferences, we used two approaches. First, in cells with genome labels, nascent transcripts were identified based on their proximity to directly observed phages. We then applied the model to the same cells and compared the phage profiles inferred by the model to those from direct genome labels, treating each profile as a clustering of mRNA spots within the cell. This comparison yielded a Rand index [32, 33] of 0.99 ± 0.06 (mean±SD, *n* = 703 cells) across all cells with genome labels (Figure S15). Second, we calculated the single-cell MOI from the inferred phage profiles and found that these values, after correction for transcriptionally silent phages (see Correcting MOI for transcriptionally silent phages), were consistent with the MOI independently determined for the same time-point samples in MOI determination by microscopy, prior to par-seqFISH (Figure 3I).

#### Correcting MOI for transcriptionally silent phages

We corrected the MOI estimates inferred by the XGBoost pipeline (see Inferring single-phage transcription profile in the absence of genome label) to account for the presence of transcriptionally silent phages. Since the number of inferred phages per cell reflects only transcriptionally active genomes, the observed value *K* may underestimate the true intracellular phage count *N*. We modeled this missingness as an independent per-phage “thinning” process [64]: each phage is silent with probability *s* and detectable with probability *q* = 1 – *s* (here *s* = 0.18, see Figure 3B–3C). Under this model, the observed count satisfies *K* | *N* ∼ Binomial(*N, q*), with likelihood

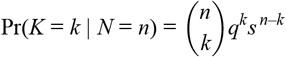

for integers *n* ≥ *k* ≥ 0. To infer the distribution of *N* from observations of *K*, we used the pre-seqFISH MOI distribution (measured from genome labels and therefore unaffected by transcriptional silence) as an empirical prior *π*(*n*) ≈ Pr(*N* = *n*), implemented as the normalized pre-seqFISH MOI histogram with a small pseudocount (1e-3) added to avoid empty bins. For each observed value *k*, we computed the posterior probability of the true MOI by Bayes’ rule [65],

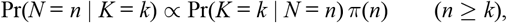

normalized over all *n* ≥ *k*. The corrected post-seqFISH MOI distribution was then obtained by redistributing the observed post-seqFISH histogram: if *c*_*k*_ denotes the number of cells with observed *K* = *k*, the expected number of cells with true MOI *n* is

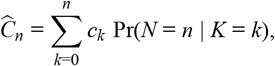

yielding 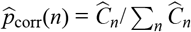 [66]. For analyses requiring a per-cell corrected MOI (e.g., calculating the mean MOI), we assigned each cell with observed *K* = *k* the posterior mean

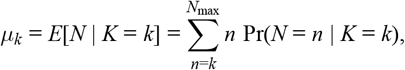

where sums were truncated at *N*_max_ chosen to exceed the maximum observed MOI by a fixed margin, set to 10 MOI units above the maximum observed value.

Uncertainty for the corrected histogram differs from that of the raw histograms because the correction distributes each observed cell fractionally over multiple true-MOI bins. Accordingly, we propagated variance from the posterior allocation rather than applying the binomial SEM used for genome labeling fractions (see Figure 3I). Treating the *c*_*k*_ cells in each observed bin *k* as independently allocating to true MOI bin *n* with probability *p*_*n*|*k*_ = Pr(*N* = *n* | *K* = *k*), the variance contribution to the corrected count in bin *n* from observed bin *k* is Var(*C*_*n*←*k*_) ≈ *c*_*k*_ *p*_*n*|*k*_(1 – *p*_*n*|*k*_), and summing across *k* gives

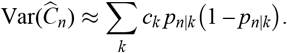

When plotting corrected fractions, we converted this to an SEM by dividing by the total number of post-seqFISH cells *C* = ∑_*k*_ *c*_*k*_, i.e.,

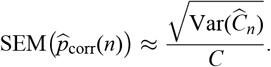

This treatment captures uncertainty introduced by ambiguity in the number of silent phages per cell under the detection model.

#### PCA on single phage data and defining phage angle

After inferring phage transcription profiles as described in Single-phage analysis, we applied the same normalization procedure used for single-cell analysis (see Principal components analysis). The normalized single-phage profiles were projected into the two-dimensional principal component space using the coefficients derived from the single-cell data (see Principal components analysis, Figure 2A). Phage angles were subsequently calculated following the approach described in Derivation of cell angle.

## Data and Code Availability

- Sequencing data have been deposited in GEO under accession numbers **GSE298101** and **GSE308589**.
- Images, codes and any additional information required to reanalyze the data reported in this paper are available from the corresponding author upon request.

## Acknowledgements

We thank L. Cai, D. Dar, Y. Geng, H. S. Han, S. Maslov, J. R. Moffitt, H. V. Nguyen, T. V. P. Nguyen, A. W. Schrader, C. M. Schroeder, M. Wang, S. Yeo, S. D. Zhao, T. Champetier and all members of the Golding lab for advice and discussions. We also thank A. G. Hernandez and D. Zheng at the Roy J. Carver Biotechnology Center at the University of Illinois at Urbana–Champaign. Work in the Golding lab is supported by the National Institutes of Health grant R35 GM140709, the National Science Foundation grant 2243257 (NSF Science and Technology Center for Quantitative Cell Biology), and the Alfred P. Sloan Foundation grant G-2023-19649.

## Author Contributions

Conceptualization: E.H., I.G.; methodology: E.H., W.Z., T.Y., I.G.; investigation: E.H.; formal analysis: E.H.; visualization: E.H.; funding acquisition: I.G.; supervision: I.G.; writing - original draft: E.H., I.G.; writing - review & editing: E.H., I.G.

## References

1. Luria, S. E. and Delbrück, M. Mutations of bacteria from virus sensitivity to virus resistance. Genetics 28 (1943), 491.

2. Delbrück, M. The burst size distribution in the growth of bacterial viruses (bacteriophages). Journal of bacteriology 50 (1945), 131–135.

3. Lieb, M. The establishment of lysogenicity in Escherichia coli. Journal of bacteriology 65 (1953), 642– 651.

4. Ptashne, M. A Genetic Switch: Phage Lambda Revisited. Cold Spring Harbor, NY: Cold Spring Harbor Laboratory Press, 2004.

5. Arkin, A., Ross, J., and McAdams, H. H. Stochastic kinetic analysis of developmental pathway bifurcation in phage λ-infected Escherichia coli cells. Genetics 149 (1998), 1633–1648.

6. Singh, A. and Weinberger, L. S. Stochastic gene expression as a molecular switch for viral latency. Current opinion in microbiology 12 (2009), 460–466.

7. Balázsi, G., Van Oudenaarden, A., and Collins, J. J. Cellular decision making and biological noise: from microbes to mammals. Cell 144 (2011), 910–925.

8. Kourilsky, P. Lysogenization by bacteriophage lambda: I. multiple infection and the lysogenic response. Molecular and General Genetics MGG 122 (1973), 183–195.

9. Zeng, L., Skinner, S. O., Zong, C., Sippy, J., Feiss, M., and Golding, I. Decision making at a sub-cellular level determines the outcome of bacteriophage infection. Cell 141 (2010), 682–691.

10. Trinh, J. T., Shao, Q., Guan, J., and Zeng, L. Emerging heterogeneous compartments by viruses in single bacterial cells. Nature Communications 11 (2020), 3813.

11. Yao, T., Coleman, S., Nguyen, T. V. P., Golding, I., and Igoshin, O. A. Bacteriophage self-counting in the presence of viral replication. Proceedings of the National Academy of Sciences 118 (2021), e2104163118.

12. Wang, B., Lin, A. E., Yuan, J., Novak, K. E., Koch, M. D., Wingreen, N. S., Adamson, B., and Gitai, Z. Single-cell massively-parallel multiplexed microbial sequencing (M3-seq) identifies rare bacterial populations and profiles phage infection. Nature microbiology 8 (2023), 1846–1862.

13. Putzeys, L., Wicke, L., Brandao, A., Boon, M., Pires, D. P., Azeredo, J., Vogel, J., Lavigne, R., and Gerovac, M. Exploring the transcriptional landscape of phage–host interactions using novel high-throughput approaches. Current Opinion in Microbiology 77 (2024), 102419.

14. Eng, C.-H. L., Lawson, M., Zhu, Q., Dries, R., Koulena, N., Takei, Y., Yun, J., Cronin, C., Karp, C., Yuan, G.-C., et al. Transcriptome-scale super-resolved imaging in tissues by RNA seqFISH+. Nature 568 (2019), 235–239.

15. Dar, D., Dar, N., Cai, L., and Newman, D. K. Spatial transcriptomics of planktonic and sessile bacterial populations at single-cell resolution. Science 373 (2021), eabi4882.

16. Gabi-Lange, D., Litvinov, V., and Dar, D. Single-Cell Phenotypic Heterogeneity Shapes Quorum Signaling Dynamics in Pseudomonas aeruginosa. BioRxiv (2025), 2025–05.

17. Wang, M., Zhang, J., Xu, H., and Golding, I. Measuring transcription at a single gene copy reveals hidden drivers of bacterial individuality. Nature microbiology 4 (2019), 2118–2127.

18. St-Pierre, F. and Endy, D. Determination of cell fate selection during phage lambda infection. Proceedings of the National Academy of Sciences 105 (2008), 20705–20710.

19. Ohno, S., Okano, H., Tanji, Y., Ohashi, A., Watanabe, K., Takai, K., and Imachi, H. A method for evaluating the host range of bacteriophages using phages fluorescently labeled with 5-ethynyl-2′-deoxyuridine (EdU). Applied microbiology and biotechnology 95 (2012), 777–788.

20. Casjens, S. R. and Hendrix, R. W. Bacteriophage lambda: Early pioneer and still relevant. Virology 479 (2015), 310–330.

21. Liu, X., Jiang, H., Gu, Z., and Roberts, J. W. High-resolution view of bacteriophage lambda gene expression by ribosome profiling. Proceedings of the National Academy of Sciences 110 (2013), 11928– 11933.

22. Skinner, S. O., Sepúlveda, L. A., Xu, H., and Golding, I. Measuring mRNA copy number in individual Escherichia coli cells using single-molecule fluorescent in situ hybridization. Nature protocols 8 (2013), 1100–1113.

23. Golding, I., Coleman, S., Nguyen, T. V., and Yao, T. Decision making by temperate phages. Encyclopedia of Virology 1 (2019).

24. Chen, H., Shiroguchi, K., Ge, H., and Xie, X. S. Genome-wide study of mRNA degradation and transcript elongation in E scherichia coli. Molecular systems biology 11 (2015), 781.

25. Silverman, A., Nashef, R., Wasserman, R., Noy, T., Born, S., Yao, T., Geng, Y., Rotbard, H., Levkowitz, A., Kaufman, Y., Golding, I., and Melamed, S. Phage-encoded small RNA hijacks host replication machinery to support the phage lytic cycle. Molecular Cell 85 (2025), 4678–4697.e12.

26. Luecken, M. D. and Theis, F. J. Current best practices in single-cell RNA-seq analysis: a tutorial. Molecular systems biology 15 (2019), e8746.

27. Pountain, A. W., Jiang, P., Yao, T., Homaee, E., Guan, Y., McDonald, K. J., Podkowik, M., Shopsin, B., Torres, V. J., Golding, I., et al. Transcription–replication interactions reveal bacterial genome regulation. Nature 626 (2024), 661–669.

28. Levin, M., Anavy, L., Cole, A. G., Winter, E., Mostov, N., Khair, S., Senderovich, N., Kovalev, E., Silver, D. H., Feder, M., et al. The mid-developmental transition and the evolution of animal body plans. Nature 531 (2016), 637–641.

29. Zalts, H. and Yanai, I. Developmental constraints shape the evolution of the nematode mid-developmental transition. Nature ecology & evolution 1 (2017), 0113.

30. Trinh, J. T., Székely, T., Shao, Q., Balázsi, G., and Zeng, L. Cell fate decisions emerge as phages cooperate or compete inside their host. Nature communications 8 (2017), 14341.

31. Chen, T. XGBoost: A Scalable Tree Boosting System. Cornell University (2016).

32. Rand, W. M. Objective criteria for the evaluation of clustering methods. Journal of the American Statistical association 66 (1971), 846–850.

33. Mirkin, B. Clustering for data mining: a data recovery approach. Chapman and Hall/CRC, 2005.

34. Zeng, L. and Golding, I. Following cell-fate in E. coli after infection by phage lambda. Journal of Visualized Experiments: JoVE (2011), 3363.

35. Weitz, J. S., Mileyko, Y., Joh, R. I., and Voit, E. O. Collective decision making in bacterial viruses. Biophysical journal 95 (2008), 2673–2680.

36. Hershey, A. D. The Bacteriophage Lambda. Ed. by Hershey, A. D. Cold Spring Harbor, NY: Cold Spring Harbor Laboratory, 1971, 792. ISBN: 9780879691028.

37. Hendrix, R. W., Roberts, J. W., Stahl, F. W., and Weisberg, R. A., eds. Lambda II. Cold Spring Harbor, New York: Cold Spring Harbor Laboratory Press, 1983. ISBN: 0-87969-150-6.

38. Isaacs, F. J., Hasty, J., Cantor, C. R., and Collins, J. J. Prediction and measurement of an autoregulatory genetic module. Proceedings of the National Academy of Sciences 100 (2003), 7714–7719.

39. Zong, C., So, L.-h., Sepúlveda, L. A., Skinner, S. O., and Golding, I. Lysogen stability is determined by the frequency of activity bursts from the fate-determining gene. Molecular systems biology 6 (2010), 440.

40. Bednarz, M., Halliday, J. A., Herman, C., and Golding, I. Revisiting bistability in the lysis/lysogeny circuit of bacteriophage lambda. PLoS one 9 (2014), e100876.

41. Oppenheim, A. B., Kobiler, O., Stavans, J., Court, D. L., and Adhya, S. Switches in bacteriophage lambda development. Annu. Rev. Genet. 39 (2005), 409–429.

42. Elowitz, M. B., Surette, M. G., Wolf, P.-E., Stock, J. B., and Leibler, S. Protein mobility in the cytoplasm of Escherichia coli. Journal of bacteriology 181 (1999), 197–203.

43. Campos, M. and Jacobs-Wagner, C. Cellular organization of the transfer of genetic information. Current opinion in microbiology 16 (2013), 171–176.

44. Ladouceur, A.-M., Parmar, B. S., Biedzinski, S., Wall, J., Tope, S. G., Cohn, D., Kim, A., Soubry, N., Reyes-Lamothe, R., and Weber, S. C. Clusters of bacterial RNA polymerase are biomolecular condensates that assemble through liquid–liquid phase separation. Proceedings of the National Academy of Sciences 117 (2020), 18540–18549.

45. Sarfatis, A., Wang, Y., Twumasi-Ankrah, N., and Moffitt, J. R. Highly multiplexed spatial transcriptomics in bacteria. Science 387 (2025), eadr0932.

46. Kuhlman, T. E. and Cox, E. C. DNA-binding-protein inhomogeneity in coli modeled as biphasic facilitated diffusion. Physical Review E (2013).

47. Echols, H., Court, D., and Green, L. On the nature of cis-acting regulatory proteins and genetic organization in bacteriophage: the example of gene Q of bacteriophage λ. Genetics 83 (1976), 5.

48. Burt, D. and Brammar, W. The cis-specificity of the Q-gene product of bacteriophage lambda. Molecular and General Genetics MGG 185 (1982), 468–472.

49. Riggs, P. D. and Botstein, D. Bacteriophage P22 gene 23 product acts preferentially in cis. Journal of virology 61 (1987), 2316–2318.

50. Golding, I. and Amir, A. Colloquium: Gene expression in growing cells: A biophysical primer. Reviews of Modern Physics 96 (2024), 041001.

51. Levesque, M. J. and Raj, A. Single-chromosome transcriptional profiling reveals chromosomal gene expression regulation. Nature methods 10 (2013), 246–248.

52. Zhang, F., Gu, W., Hurles, M. E., and Lupski, J. R. Copy number variation in human health, disease, and evolution. Annual review of genomics and human genetics 10 (2009), 451–481.

53. Sambrook, J. and Russell, D. W. Molecular Cloning: A Laboratory Manual. 3rd ed. Cold Spring Harbor, NY: Cold Spring Harbor Laboratory Press, 2001.

54. Hu, M., Yang, B., Cheng, Y., Radda, J. S., Chen, Y., Liu, M., and Wang, S. ProbeDealer is a convenient tool for designing probes for highly multiplexed fluorescence in situ hybridization. Scientific Reports 10 (2020), 22031.

55. Altschul, S. F., Gish, W., Miller, W., Myers, E. W., and Lipman, D. J. Basic local alignment search tool. Journal of molecular biology 215 (1990), 403–410.

56. Reichardt, L. F. Control of bacteriophage lambda repressor synthesis after phage infection: the role of the N, cII, cIII and cro products. Journal of molecular biology 93 (1975), 267–288.

57. Geng, Y., Nguyen, T. V. P., Homaee, E., and Golding, I. Using bacterial population dynamics to count phages and their lysogens. Nature communications 15 (2024), 7814.

58. Davison, A. C. and Hinkley, D. V. Bootstrap methods and their application. Cambridge university press, 1997.

59. Nguyen, T. V. P., Wu, Y., Yao, T., Trinh, J. T., Zeng, L., Chemla, Y. R., and Golding, I. Coinfecting phages impede each other’s entry into the cell. Current Biology 34 (2024), 2841–2853.

60. Application Note 08: Coating Protocols for ibidi Labware. Version 6.2. Accessed 15 December 2025. ibidi GmbH. Gräfelfing, Germany, 2025.

61. Ronneberger, O., Fischer, P., and Brox, T. “U-net: Convolutional networks for biomedical image segmentation”. International Conference on Medical image computing and computer-assisted intervention. Springer. 2015, 234–241.

62. O’Connor, O. M., Alnahhas, R. N., Lugagne, J.-B., and Dunlop, M. J. DeLTA 2.0: A deep learning pipeline for quantifying single-cell spatial and temporal dynamics. PLoS computational biology 18 (2022), e1009797.

63. Bishop, C. M. and Nasrabadi, N. M. Pattern recognition and machine learning. Vol. 4. Springer, 2006.

64. Daley, D. J. and Vere-Jones, D. An introduction to the theory of point processes: volume I: elementary theory and methods. Springer, 2003.

65. Reich, B. J. and Ghosh, S. K. Bayesian statistical methods. Chapman and Hall/CRC, 2019.

66. Gelman, A., Carlin, J. B., Stern, H. S., and Rubin, D. B. Bayesian data analysis. Chapman and Hall/CRC, 1995.

