## Supplementary Material for "Single-phage profiling illuminates viral individuality during cell fate determination"

*This file includes:*

### SUPPLEMENTARY FIGURES

S1 Single-cell MOI distribution measured by ParB-*parS* labeling and comparison to EdU-Click. . . . . . . 2
S2 The transcription kinetics of $\lambda$ genes across biological replicates. . . . . . . 3
S3 Promoter activity kinetics during infection by *P*+ and *P*– phages. . . . . . . 4
S4 Inferring the elongation speed of RNAP from transcription progression along the $\lambda$ genome. . . . . . . 5
S5 Reproducibility of par-seqFISH measurements and agreement with bulk RNA-seq during infection by $\lambda_{\mathrm{TY5}}$ (*P*+). . . . . . . 6
S6 Gene expression kinetics during infection by $\lambda_{\mathrm{TY5}}$ (*P*+) and $\lambda_{\mathrm{TY11}}$ (*P*–) phages. . . . . . . 7
S7 Captured variance by principal components. . . . . . . 8
S8 Reproducibility of the principal-component loadings across biological replicates. . . . . . . 9
S9 Promoter activity in the PC1–PC2 plane. . . . . . . 10
S10 Reproducibility of par-seqFISH expression measurements and agreement with bulk RNA-seq during infection by $\lambda_{\mathrm{TY11}}$ (*P*–) phage. . . . . . . 11
S11 Principal-component analysis of cell fate during infection by $\lambda_{\mathrm{TY11}}$ (*P*–). . . . . . . 12
S12 The distribution of distances between mRNA spots and the nearest EdU-Click-labeled phage genome. . . . 13
S13 Genome-label deterioration during par-seqFISH reduces the number of detected phage genomes. . . . . 14
S14 Pairwise mRNA distance distributions. . . . . . . 15
S15 Patch-level mRNA copy number distributions and agreement between genome-based and model-based phage profiles. . . . . . . 16
S16 Single-phage transcription profiles and their principal-component structure. . . . . . . 17
S17 The distributions of $\theta_{\mathrm{phage}} - \theta_{\mathrm{cell}}$ for cell–phage pairs in the different developmental stages. . . . . . . 18

### SUPPLEMENTARY TABLES

S1 Bacterial strains, phages, and plasmids used in this study. . . . . . . 19
S2 Growth media and buffers used in this study. . . . . . . 20
S3 Chemicals, reagents, and kits used in this study. . . . . . . 21
S4 Gene library used in this study. . . . . . . 23
S5 16S rRNA time-point tagging probe sequences. . . . . . . 26

### SUPPLEMENTARY FIGURES

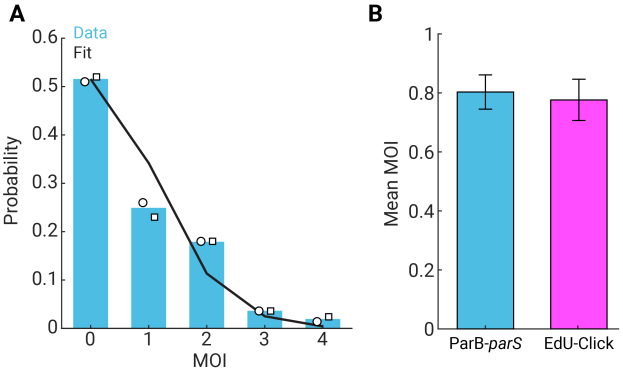


**Figure S1. Single-cell MOI distribution measured by ParB-*parS* labeling and comparison to EdU-Click.**

(A) The distribution of multiplicity of infection (MOI) in individual cells measured by ParB-*parS* labeling. Circles and squares show data from $t$ = 5 min ($n$ = 138 cells) and $t$ = 20 min ($n$ = 165 cells), respectively. Black line, fit to a Poisson distribution (with an inferred mean of 0.66).

(B) The mean MOI values obtained from ParB-*parS* and EdU-Click genome labeling. For each method, the mean was computed from the combined $t$ = 5 and $t$ = 20 min data. The total number of analyzed cells was $n$ = 303 for ParB-*parS* and $n$ = 320 for EdU–Click. Error bars report the standard error of the mean (SEM) across cells.

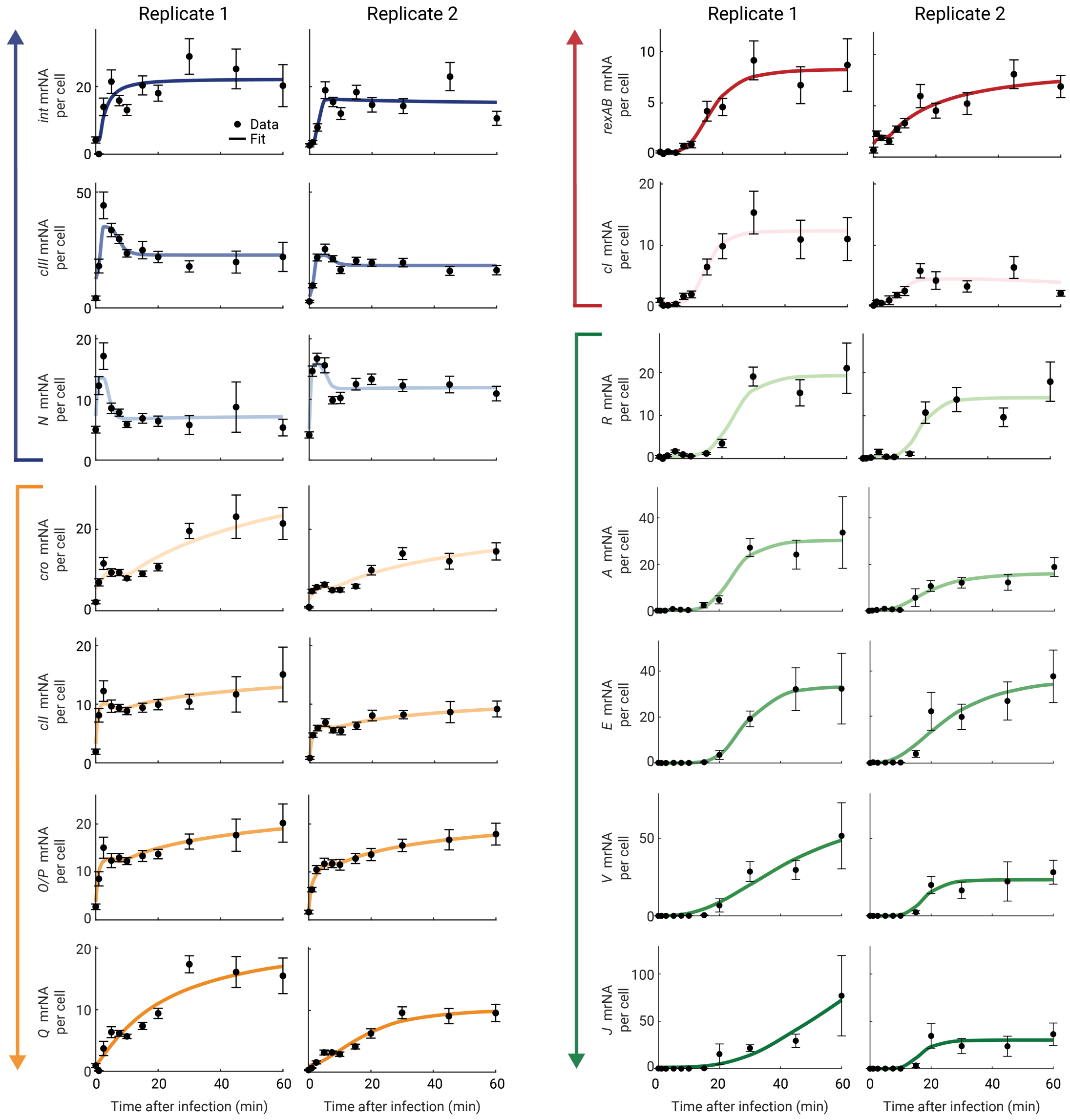


**Figure S2. The transcription kinetics of $\lambda$ genes across biological replicates.**
The mRNA copy number per cell of *int*, *cIII*, *N*, *cro*, *cII*, $\{O, P\}$, $Q$, $\{rexA, rexB\}$, *cI*, *R*, *A*, *E*, *V*, and *J*, at different times following infection of MG1655 by $\lambda_{\text{TY5}}$ ($P$+), shown for two biological replicates. Arrows on the left indicate the corresponding operons. Points, the mean across cells at each time point (replicate 1: $n = 31-160$ infected cells per time point; replicate 2: $n = 90-262$ infected cells per time point); error bars, SEM; Solid lines, fits to a sum of two Hill functions, used as a guide to the eye.

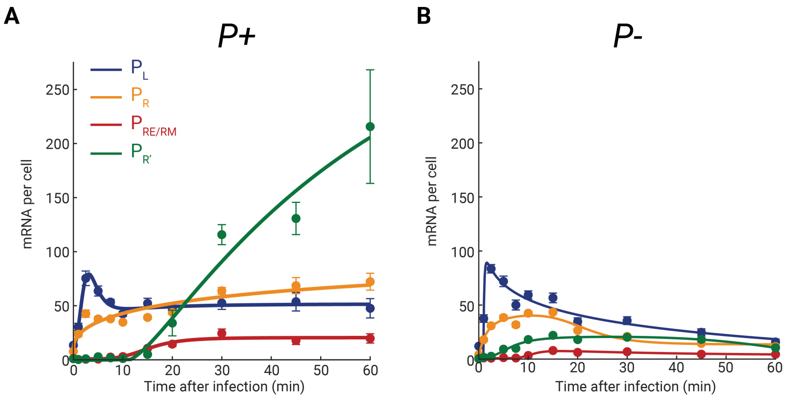


**Figure S3. Promoter activity kinetics during infection by *P*+ and *P*– phages.**
(A) Promoter activity following infection by $\lambda_{\mathrm{TY5}}$ (*P*+), quantified as the copy number per cell of transcripts assigned to each promoter, shown as a function of time after infection. Points, the mean across cells at each time point ($n$ = 31 – 160 infected cells per time point); error bars, SEM; Solid lines, fits to a sum of two Hill functions, used as a guide to the eye.
(B) Promoter activity following infection by $\lambda_{\mathrm{TY11}}$ (*P*–), calculated and plotted as in (A) ($n$ = 94 – 312 infected cells per time point).

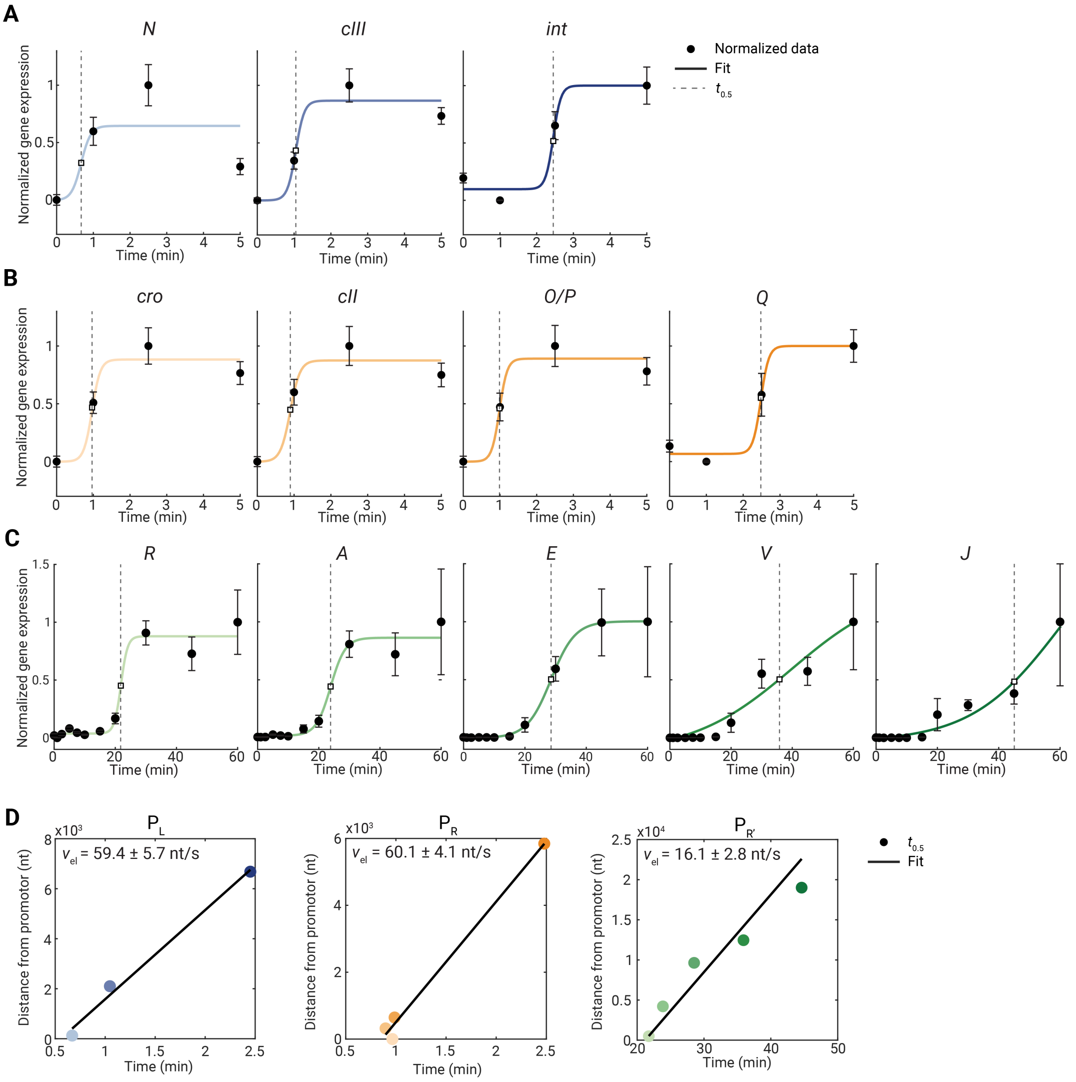


**Figure S4. Inferring the elongation speed of RNAP from transcription progression along the $\lambda$ genome.**

(A) Normalized expression trajectories for genes transcribed from the $P_L$ promoter (*N*, *cIII*, *int*) during infection by $\lambda_{TY5}$ (*P*+), over an early-time window (truncated before the onset of repression). Points, the mean expression across all cells at each time point, normalized by subtracting the minimum and dividing by the maximum across time; $n$ = 31 – 160 infected cells per time point; error bars, SEM. Solid curves, fit to a sigmoid function. Vertical dashed lines mark $t_{0.5}$, defined as the time at which the fitted trajectory reaches the midpoint between its fitted minimum and maximum values.

(B) Normalized expression trajectories for $P_R$ genes (*cro*, *cII*, *O*/*P*, *Q*), quantified and plotted as in (A).

(C) Normalized expression trajectories for $P_{R'}$ genes (*R*, *A*, *E*, *V*, *J*), quantified and plotted as in (A).

(D) The genomic distance from the promoter to each gene is plotted versus $t_{0.5}$ for each operon, and the elongation speed of RNAP ($v_{el}$) is obtained from the slope of a linear regression; reported uncertainties denote the standard error of the regression slope.

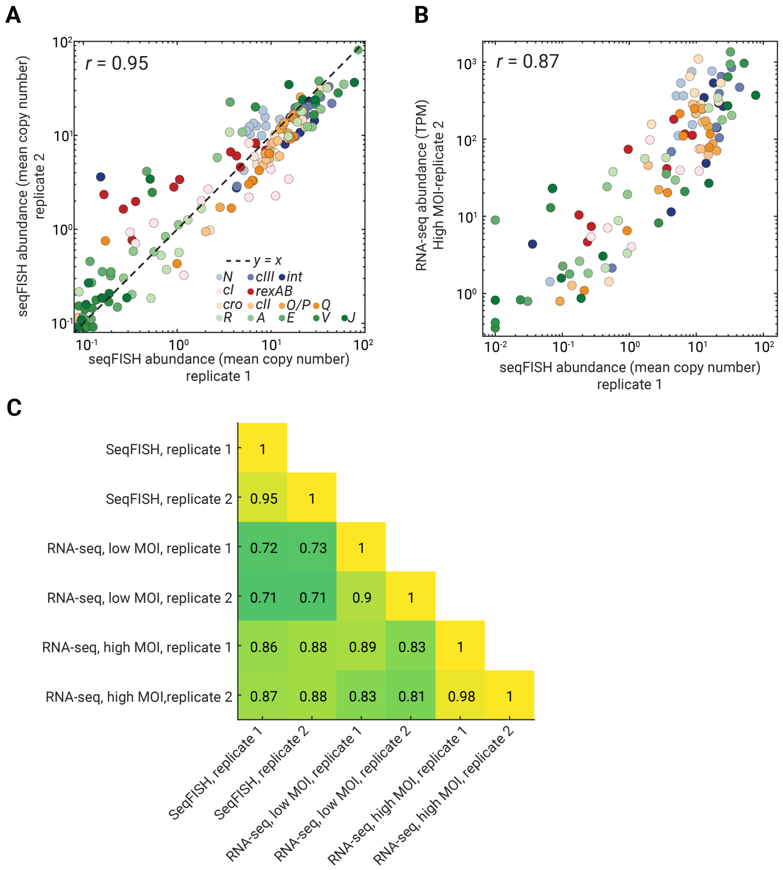


**Figure S5. Reproducibility of par-seqFISH measurements and agreement with bulk RNA-seq during infection by $\lambda_{TY5}$ (*P*+).**
(A) The average mRNA copy number per cell determined for replicate 1 versus replicate 2 of par-seqFISH for *P*+ infection. Each point indicates the mean expression of one gene at a single time point (or a control sample). Colors indicate gene identity. *r* denotes the Pearson correlation coefficient between the logarithmic expression values.
(B) The average mRNA copy number per cell determined by par-seqFISH (replicate 1) versus the RNA abundance determined by bulk RNA-seq (transcripts per million reads, TPM) for the same genes at matched time points for *P*+ infection. Each point indicates the mean expression value of one gene at a single time point (or a control sample). Colors indicate gene identity. *r* denotes the Pearson correlation coefficient between the logarithmic expression values.
(C) All pairwise correlation coefficients between the logarithmic average mRNA copy number per cell determined for all combinations of par-seqFISH replicates and bulk RNA-seq conditions during infection by *P*+ phage.

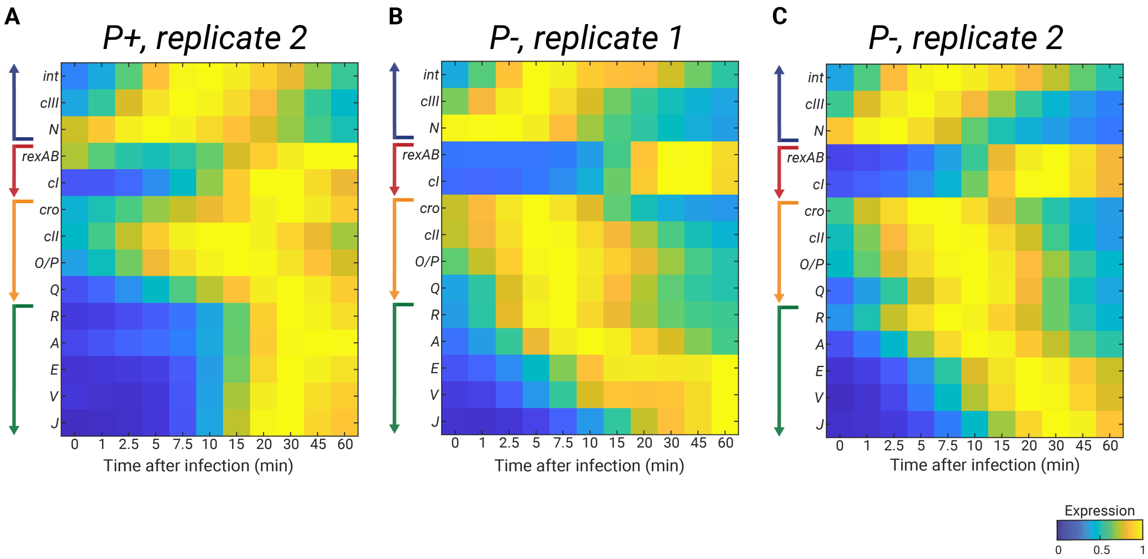


**Figure S6. Gene expression kinetics during infection by $\lambda_{TY5}$ (*P*+) and $\lambda_{TY11}$ (*P*–) phages.**
(A) Expression kinetics of $\lambda$ genes following infection by *P*+ phage (replicate 2; the first replicate is shown in Figure 1F). The cell-averaged mRNA copy number for each gene, as a function of time ($n$ = 90 – 262 infected cells per time point) was smoothed by fitting to a sum of two Hill functions, and the resulting values were normalized by the maximal value across all samples. Arrows on the left indicate the corresponding phage operons.
(B) Similar to (**A**), infection by *P*– phage (replicate 1); $n$ = 56 – 125 infected cells per time point.
(C) Similar to (**A**), infection by *P*– phage (replicate 2); $n$ = 94 – 312 infected cells per time point.

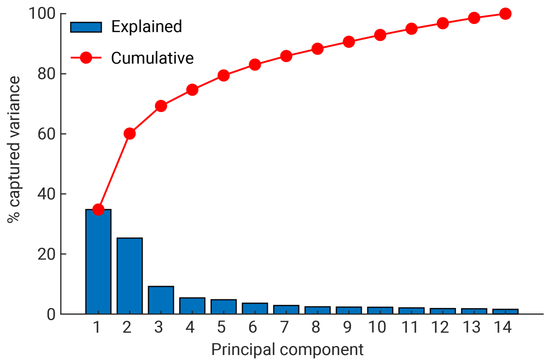


**Figure S7. Captured variance by principal components.**
PCA was applied to par-seqFISH gene-expression data from the 14 targeted $\lambda$ genes across all samples during infection by $\lambda_{TY5}$ (*P*+), as described in METHODS. Bars report the fraction of variance explained by each principal component. The line reports the cumulative variance explained.

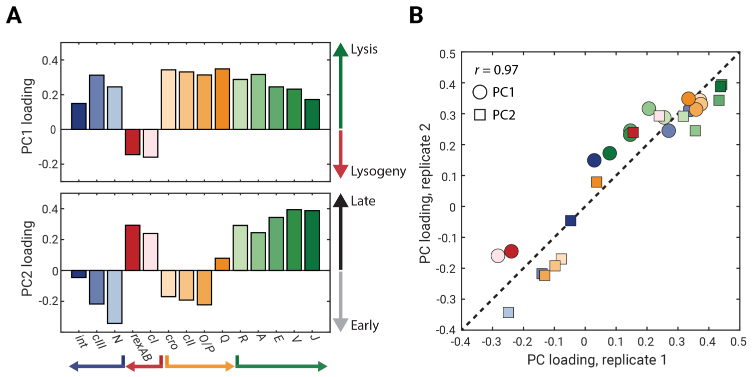


**Figure S8. Reproducibility of the principal-component loadings across biological replicates.**
(A) PCA eigenvectors of the single-cell gene expression data. The loadings of the first two principal component vectors derived from the second biological replicate of the $\lambda_{TY5}$ ($P$+) infection dataset ($n$ = 2187 cells; the first replicate is shown in Figure 2A). $\lambda$ genes and operons are colored as in Figure 1D. Arrows on the right indicate the correspondence with lysis-vs.-lysogeny (PC1, top) and early-vs.-late (PC2, bottom).
(B) The PC loadings for replicate 1 versus replicate 2 for $P$+ infection. Points report gene loadings for PC1 (circles) and PC2 (squares), with point colors indicating gene identity. The dashed line indicates $y = x$. $r$ denotes the Pearson correlation coefficient between replicate loadings.

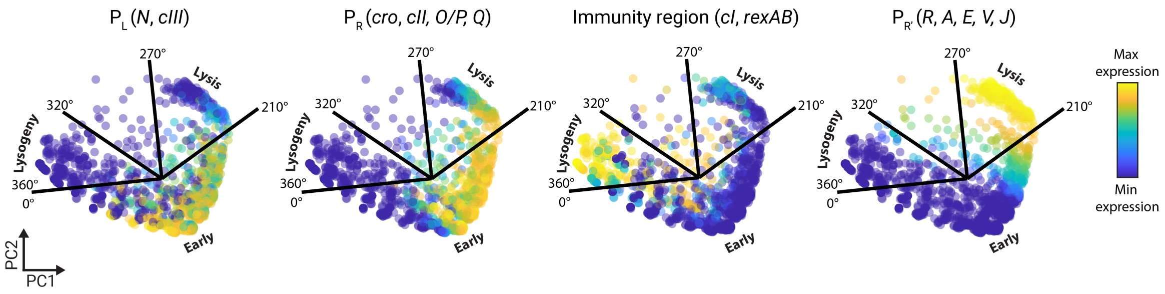


**Figure S9. Promoter activity in the PC1–PC2 plane.**
Promoter activity in the PC1–PC2 plane during infection by $\lambda_{\mathrm{TY5}}$ (*P*+), shown by coloring cells according to the natural log of normalized promoter-level expression (quantified as in Figure S3) . Radial boundaries define sectors corresponding to early, lytic, and lysogenic regions, as indicated.

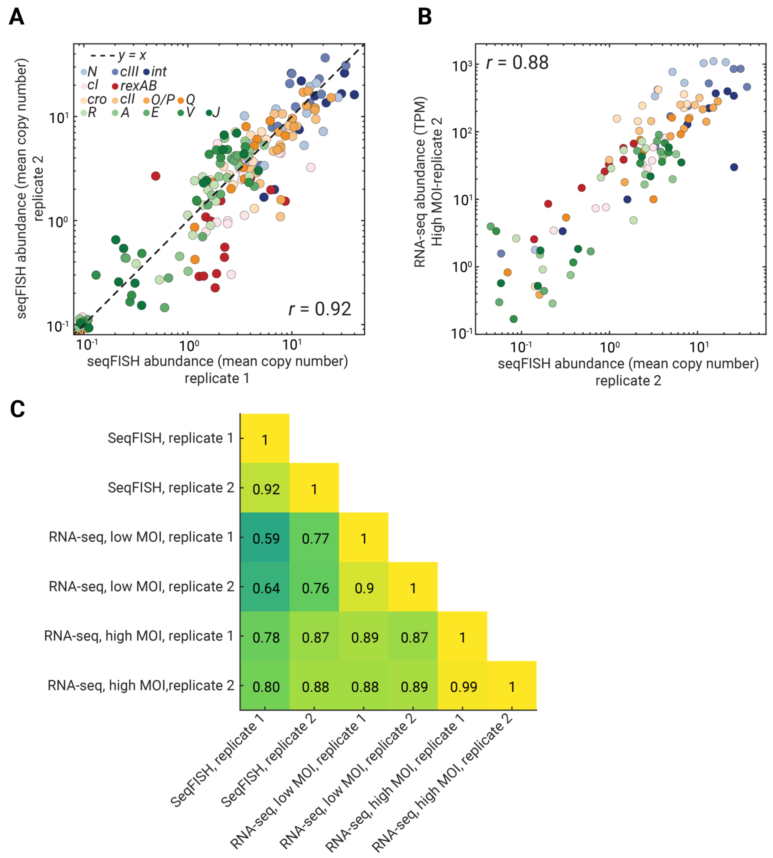


**Figure S10. Reproducibility of par-seqFISH expression measurements and agreement with bulk RNA-seq during infection by $\lambda_{TY11}$ (*P*–) phage.**

(A) The average mRNA copy number per cell determined for replicate 1 versus replicate 2 of par-seqFISH for *P*– infection. Each point indicates the mean expression of one gene corresponding to a single time point (or a control sample). Colors indicate gene identity. *r* denotes the Pearson correlation coefficient between the logarithmic expression values.

(B) The average mRNA copy number per cell determined by par-seqFISH (replicate 2) versus the RNA abundance determined by bulk RNA-seq (TPM) for the same genes at matched time points for *P*– infection. Each point indicates the mean expression value of one gene corresponding to a single time point (or a control sample). Colors indicate gene identity. *r* denotes the Pearson correlation coefficient between the logarithmic expression values.

(C) All pairwise correlation coefficients between the logarithmic average mRNA copy number per cell determined for all combinations of par-seqFISH replicates and bulk RNA-seq conditions during infection by *P*– phage.

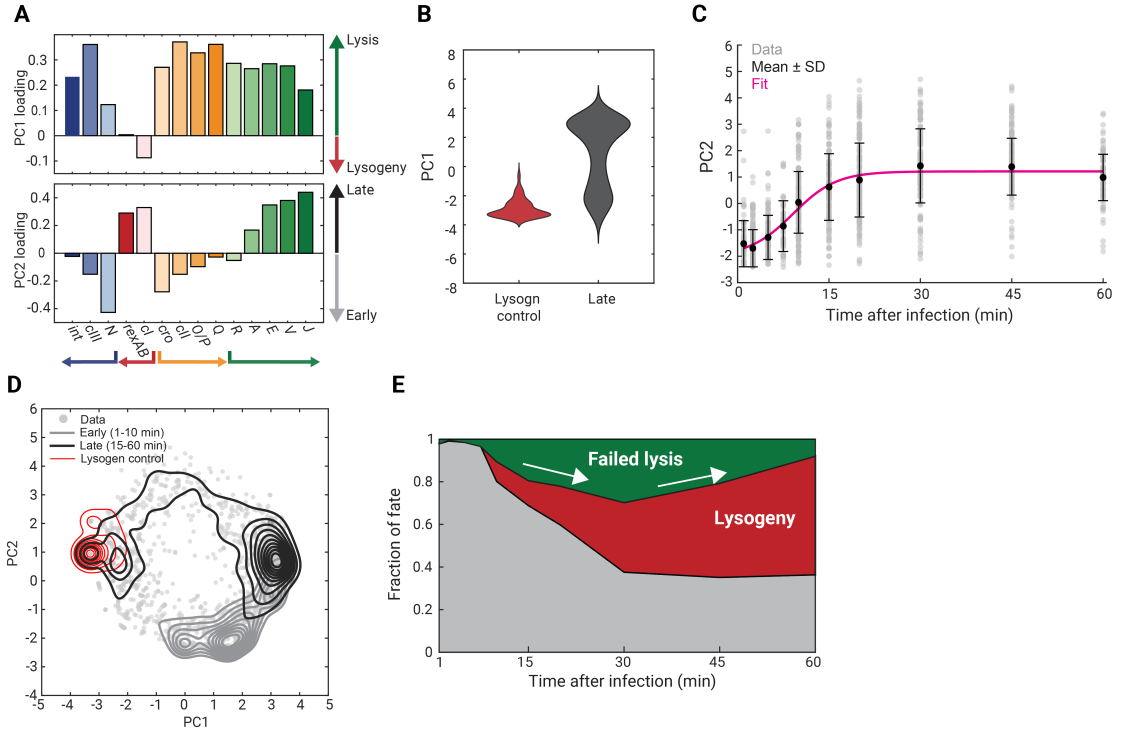


**Figure S11. Principal-component analysis of cell fate during infection by $\lambda_{\mathrm{TY11}}$ (*P*–).**
(A) PCA eigenvectors of the single-cell gene expression data. The coefficients (loadings) of the first two principal component vectors derived from the *P*– single-cell RNA dataset ($n$ = 2,203 cells). $\lambda$ genes and operons are colored as in Figure 1D. Arrows on the right indicate the correspondence with lysis-vs.-lysogeny (PC1, top) and early-vs.-late (PC2, bottom).
(B) Interpretation of PC1 for *P*– infection. PC1 scores for lysogen control ($n$ = 104 cells) and for late-stage infected cells (pooled from $t = 15, 20, 30, 45, 60$ min; $n$ = 888), where both lytic (PC1$>$ 0) and lysogenic (PC1$<$ 0) populations are visible.
(C) Interpretation of PC2 for *P*– infection. PC2 scores as a function of time after infection (94 – 312 cells per sample), demonstrating the monotonic increase with time. Gray points, individual cells; black markers, mean $\pm$ SD at each time point; pink line, fit to a sigmoid function, serving as a guide to the eye.
(D) Infected cell states in the PC1–PC2 plane. The expression profile of each cell was projected to the plane of PC1 and PC2. Gray point, individual cells; solid lines, kernel-density contours for early-stage (1–10 min, light gray, $n$ = 779), late stage (15–60 min, dark gray, $n$ = 888), and lysogen control cells (red, $n$ = 104).
(E) The emergence of cell fates during *P*– infection. Based on the angular regions defined by $\theta_{\mathrm{cell}}$ (early: $0^\circ$ – $210^\circ$; failed lytic: $210^\circ$ – $270^\circ$; lysogenic: $270^\circ$ – $360^\circ$), cells at each time point were scored as lytic (red), lysogenic (green), or early (gray). The colored areas indicate the fraction of cells in each category over time.

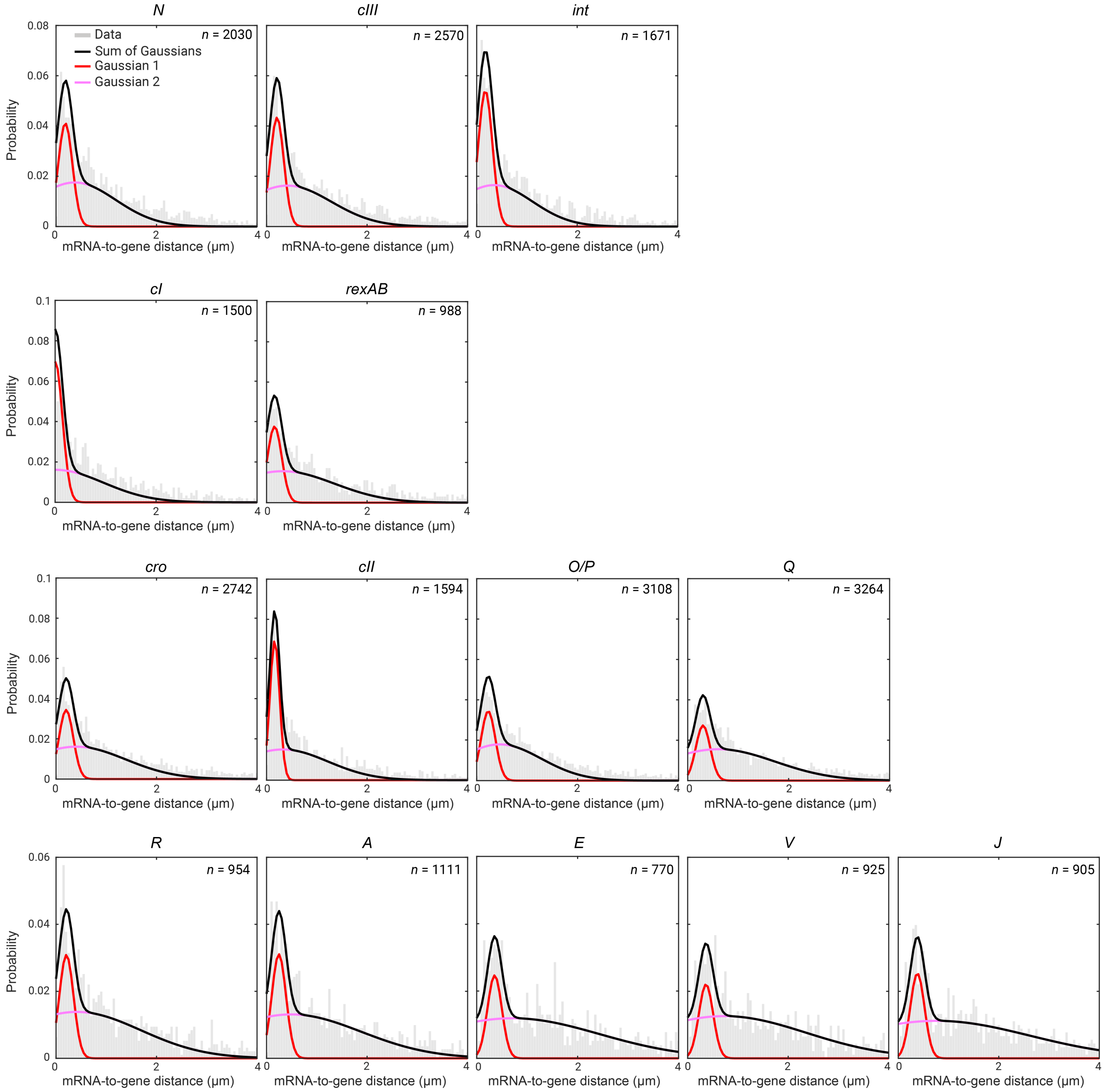


**Figure S12. The distribution of distances between mRNA spots and the nearest EdU-Click-labeled phage genome.**
For each mRNA species, the distance from each mRNA spot to its nearest phage genome (EdU-Click labeled) position within the same cell was computed. Histograms show the distribution of mRNA-to-genome distances during infection by $\lambda_{\mathrm{TY5}}$ (*P*+); the number of mRNA–genome pairs is reported in each panel. The distributions were fitted to a sum of two Gaussians, representing a genome-proximal population and a more distal population. mRNAs falling within the first Gaussian, centered near the genome, were classified as nascent, while those in the second Gaussian were classified as mature.

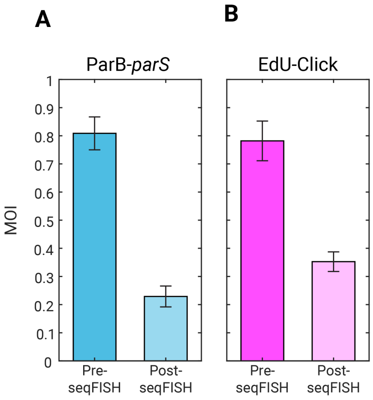


**Figure S13. Genome-label deterioration during par-seqFISH reduces the number of detected phage genomes.**

(A) Mean MOI values obtained from ParB-*parS* genome labeling before and after par-seqFISH during infection by $\lambda_{\mathrm{TY5}}$ (*P*+) (pooled from $t$ = 5, 20,min; pre-seqFISH, $n$ = 303 cells; post-seqFISH, $n$ = 485 cells). Bars, mean; error bars, SEM.

(B) Mean MOI values obtained from EdU-Click genome labeling before and after par-seqFISH during infection by $\lambda_{\mathrm{TY5}}$ (*P*+) (pooled from $t$ = 5, 20,min; pre-seqFISH, $n$ = 320 cells; post-seqFISH, $n$ = 485 cells). Bars, mean; error bars, SEM.

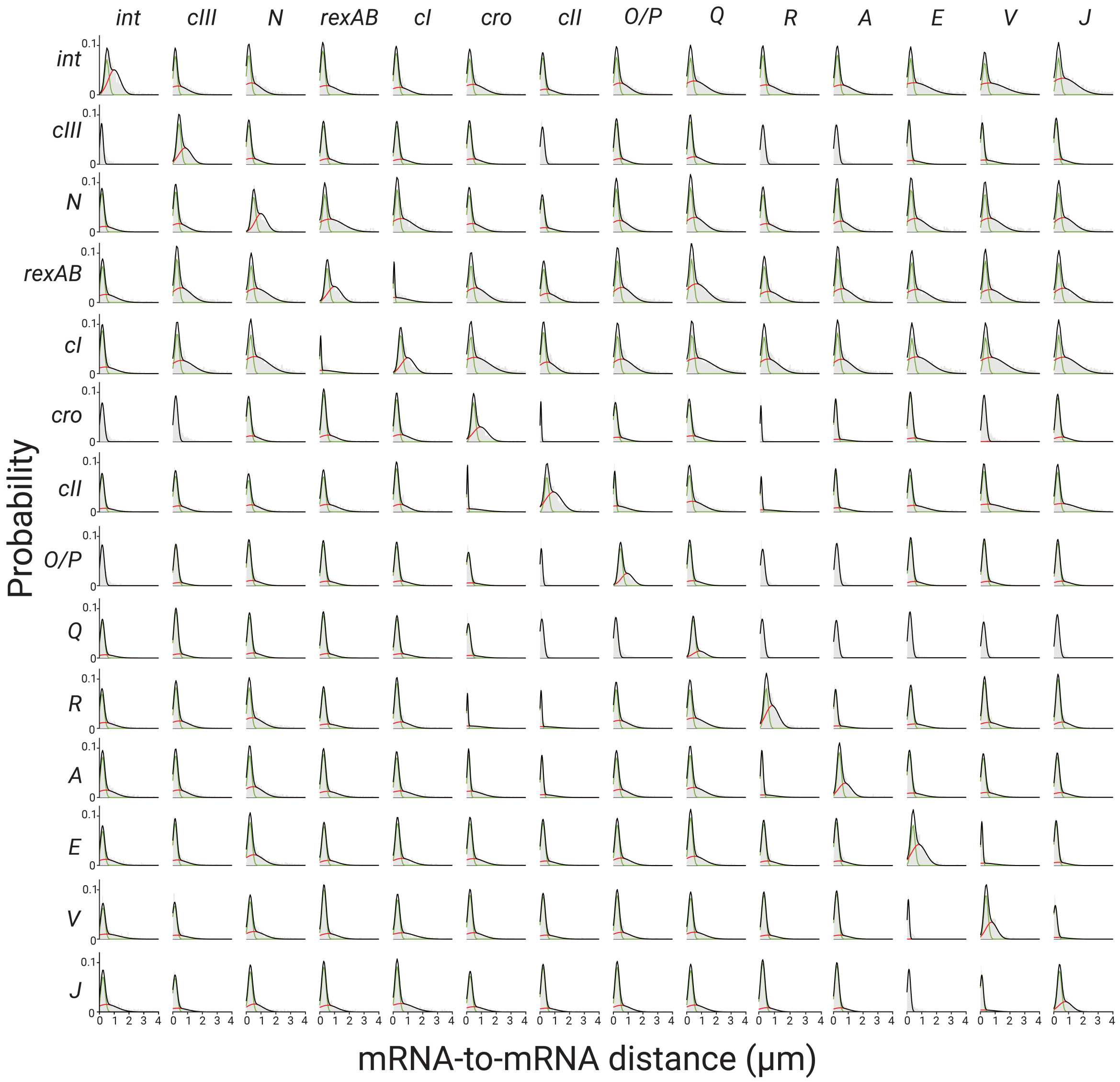


**Figure S14. Pairwise mRNA distance distributions.**
The distribution of distances between nearest mRNA molecules of different species during infection by $\lambda_{TY5}$ (*P*+), shown for all gene pairs. For each pair, the distance from each mRNA spot of one species to the nearest mRNA spot of the other species within the same cell was computed. Histograms show the resulting mRNA–mRNA distance distributions; per gene pair, *n* = 723–6,704 pairs from *n* = 257 – 1,410 cells. Black, fits to a sum of two Gaussians, corresponding to a colocalized (proximal) population and a more distal population. Bin widths were varied across gene pairs to improve visibility of the distributions.

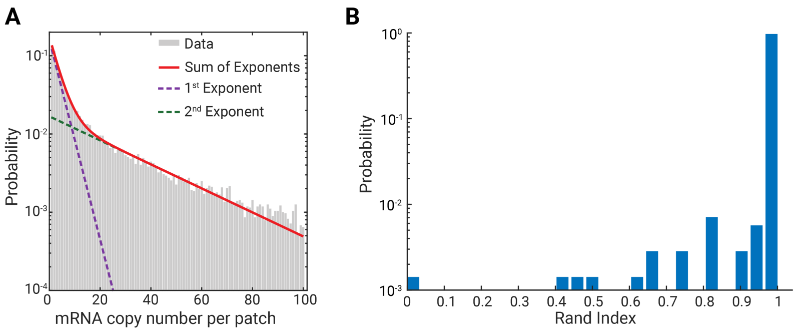


**Figure S15. Patch-level mRNA copy number distributions and agreement between genome-based and model-based phage profiles.**
(A) The distribution of total mRNA copy number per patch during infection by $\lambda_{TY5}$ ($P$+). Red, fit to a sum of two exponential functions, separating a low-count regime (interpreted as the random colocalization of mature mRNA) and a higher-count regime (interpreted as the colocalization of nascent mRNA at active transcription sites) (see METHODS). Dashed lines, individual exponential components.
(B) The distribution of Rand index values comparing genome-based and model-based phage profiles. In cells with detected phage genomes, nascent transcripts were identified based on their proximity to labeled phage genomes, and mRNA spots were grouped into individual phage profiles (see METHODS). The trained model was then applied to these detected nascent transcripts (see METHODS) and the phage profiles inferred by the model were compared to those from direct genome labels, treating each profile as a cluster of mRNA spots within the cell ($n$ = 703 cells; mean $\pm$ SD = 0.99 $\pm$ 0.06).

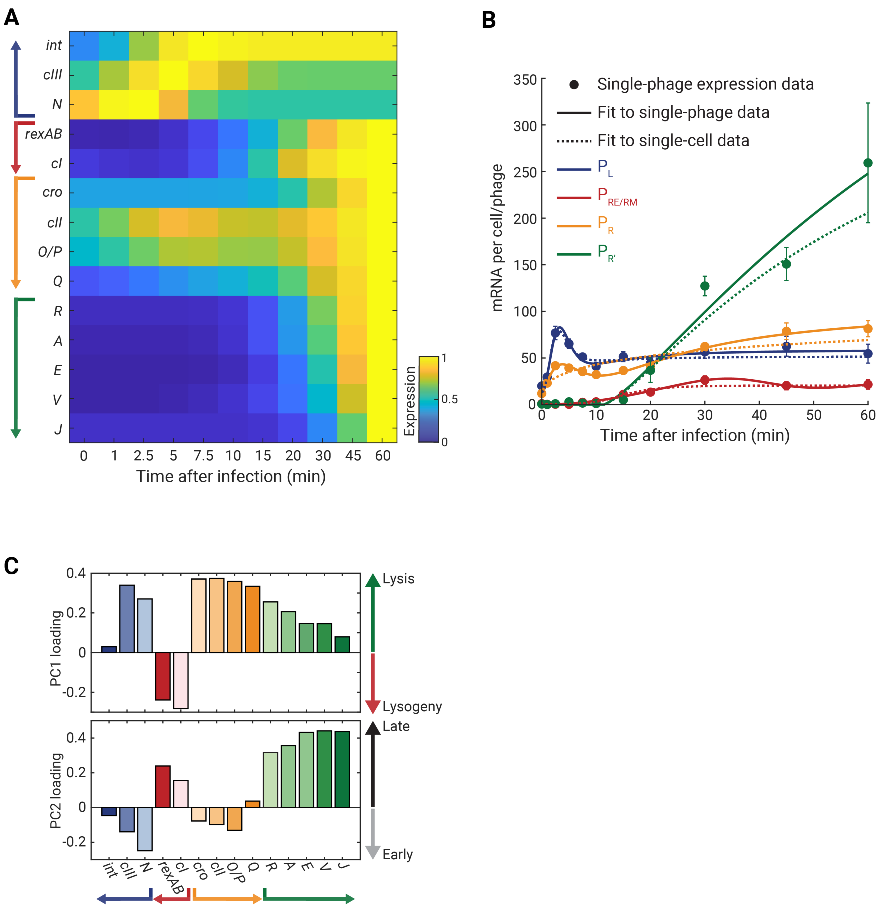


**Figure S16. Single-phage transcription profiles and their principal-component structure.**
(A) Single-phage expression kinetics during infection by $\lambda_{TY5}$ ($P$+). The phage-averaged mRNA copy number for each gene, as a function of time ($n$ = 67 – 228 phages per time point) was smoothed by fitting to a sum of two Hill functions, and the resulting values were normalized by the maximal value across all samples.
(B) Promoter activity over time for single-phage transcription profiles (solid lines) compared to the activity computed from single-cell data (dotted lines). The single-cell fits are taken from Figure S3. Promoter activity was quantified as in Figure S3. Points report the mean across phages at each time point; error bars report SEM ($n$ = 67 – 228 phages per time point).
(C) PCA eigenvectors of the single-phage gene expression data. The coefficients (loadings) of the first two principal component vectors derived from the $P$+ single-phage RNA dataset ($n$ = 2,586 phages). $\lambda$ genes and operons are colored as in Figure 1D. Arrows on the right indicate the correspondence with lysis-vs.-lysogeny (PC1, top) and early-vs.-late (PC2, bottom).

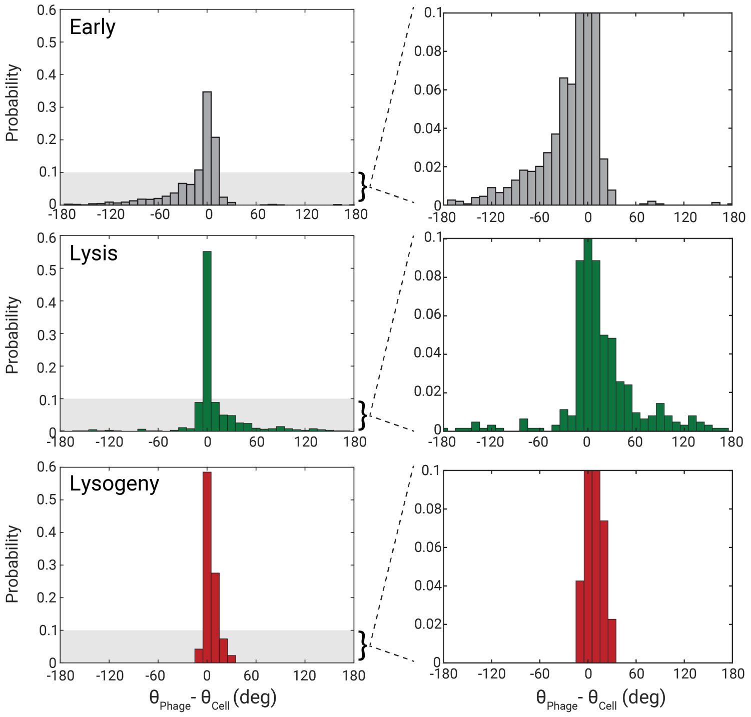


**Figure S17. The distributions of $\theta_{\text{phage}} - \theta_{\text{cell}}$ for cell–phage pairs in the different developmental stages.**
Data from $n$ = 1,541 early, $n$ = 620 lytic, and $n$ = 352 lysogenic phage-cell pairs. Right panels show magnified views of the corresponding distributions.

### Supplemental Tables

**Table S1. Bacterial strains, phages, and plasmids used in this study.**

| Bacterial strains | | |
|---|---|---|
| **Strain name** | **Relevant genotype or description** | **Source** |
| MG1655 | Wild-type | Lab stock |
| LE392 | *glnV* (*supE*44), *tryT* (*supF*58) | Lab stock |
| MG1655 $\lambda_{TY5}$ | $\lambda$ *cI857 Sam7 stf*::P1*parS-kan*$^{R}$ | Lab stock |
| MG1655 $\lambda_{TY9}$ | $\lambda$ *cI857 stf*::P1*parS-kan*$^{R}$ | Lab stock |
| MG1655 $\lambda_{TY11}$ | $\lambda$ *cI857 Pam80 stf*::P1*parS-kan*$^{R}$ | Lab stock |
| **Phage strains** | | |
| **Strain name** | **Relevant genotype or description** | **Source** |
| $\lambda_{TY5}$ | $\lambda$ *cI857 Sam7 stf*::P1*parS-kan*$^{R}$ | Lab stock |
| $\lambda_{TY11}$ | $\lambda$ *cI857 Pam80 stf*::P1*parS-kan*$^{R}$ | Lab stock |
| **Plasmids** | | |
| **Plasmid name** | **Description** | **Source** |
| pALA3047 | $P_{lac}$-*cfp*-P1-$\Delta$30*parB*-*amp*$^{R}$ | Stuart Austin |

**Table S2. Growth media and buffers used in this study.**

| Medium or buffer | Recipe | Storage condition & Notes |
|---|---|---|
| LB (Lennox) | By w/v, 1% tryptone, 0.5% yeast extract, and 0.5% NaCl; pH adjusted using 1 mM NaOH. | RT* after autoclaving. |
| LBGM | LB, 0.2% glucose, and 10 mM $MgSO_4$. | Made fresh for each experiment. |
| LBMM | LB, 0.2% maltose, and 10 mM $MgSO_4$. | Made fresh for each experiment. |
| NZYM | By w/v, 2.2% NZYM medium; pH adjusted using 10 mM NaOH. | 4°C after autoclaving. |
| SM | 100 mM NaCl, 10 mM $MgSO_4$, 50 mM Tris-Cl (pH 7.5), and 0.01% (w/v) gelatin. | RT. |
| PBS | 1×, prepared from a 10× stock. | RT. |
| SSC | 2× and 4×, prepared from a 20× stock. | RT. |
| TE | 10 mM Tris-HCl (pH 8.0), 0.1 mM EDTA. | RT. |

* RT, room temperature.

**Table S3.** **Chemicals, reagents, and kits used in this study.**

| Reagent | Supplier | Cat. # | Working concentration |
|---|---|---|---|
| Ampicillin | Fisher Scientific | BP1760-25 | 100 mg/ml |
| Kanamycin | Fisher Scientific | BP906-5 | 50 mg/ml |
| NaCl | Fisher Scientific | BP358-1 | 5 M |
| NaOH | Fisher Scientific | BP359-500 | 1 M |
| $MgSO_4$ | Fisher Scientific | BP213-1 | 1 M |
| Maltose | Fisher Scientific | BP684-500 | 20% |
| Glucose | Fisher Scientific | BP350-500 | 20% |
| Isopropyl-$\beta$-thiogalactoside (IPTG) | Sigma-Aldrich | I6758 | 500 mM |
| Formaldehyde | Fisher Scientific | BP531-500 | 37% |
| Chloroform | Fisher Scientific | C298-500 | — |
| DEPC-treated water | Fisher Scientific | 4387937 | — |
| Agarose | Sigma-Aldrich | A9414 | — |
| Agar | BD Biosciences | 214010 | — |
| NZYM | Teknova | N2062 | — |
| Tryptone | Gibco | 211705 | — |
| Yeast extract | Gibco | 212750 | — |
| Ethanol | Decon Labs | 2716 | — |
| 0.01% Poly-L-Lysine | Sigma-Aldrich | P4707 | 0.01% |
| 10× PBS | Fisher Scientific | AM9624 | 10× |
| 20× SSC | Fisher Scientific | AM9763 | 20× |
| Formamide | Fisher Scientific | AM9342 | — |
| Dextran sulfate | Sigma-Aldrich | D8906 | — |
| Triton X-100 | Sigma-Aldrich | 93443 | — |

**Table S3.** **Table S3 (continued)**

| Reagent | Supplier | Cat. # | Working concentration |
|---|---|---|---|
| Ethylene carbonate | Sigma-Aldrich | E26258 | — |
| TE buffer | Fisher Scientific | BP2473100 | — |
| DNase I | Zymo | E1010 | — |
| ibidi $\mu$-Slide VI 0.5 | ibidi | 80607 | — |
| RNAprotect Bacteria Reagent | Qiagen | 76506 | — |
| Quick-RNA Fungal/Bacterial Microprep Kit | Zymo Research | R2010 | — |
| Click-iT EdU Cell Proliferation Kit for Imaging, Alexa Fluor™ 647 dye | Fisher Scientific | C10340 | — |

**Table S4.** **Gene library used in this study.**

| Gene | mRNA binding region sequence | Full primary probe sequence |
|---|---|---|
| *int* | CCGGCGCTCATGACTTCGCCTTCTTCCCAT | TGATACGCGAACTGTAATGATACGCGAACTGTAACCGGCGCTCATGACTTCGCCTTCTTCCCATAATGATACGCGAACTGTAATGATACGCGAACTGT |
| *int* | CTTGGGTCCCTGTAGCAGTAATATCCATTG | TGATACGCGAACTGTAATGATACGCGAACTGTAACTTGGGTCCCTGTAGCAGTAATATCCATTGAATGATACGCGAACTGTAATGATACGCGAACTGT |
| *int* | CCTGTATAGCTTCAGTGATTGCGATTCGCC | TGATACGCGAACTGTAATGATACGCGAACTGTAACCTGTATAGCTTCAGTGATTGCGATTCGCCAATGATACGCGAACTGTAATGATACGCGAACTGT |
| *int* | CTGTCAGAGGCTTGTGTTTGTGTCCTGAAA | TGATACGCGAACTGTAATGATACGCGAACTGTAACTGTCAGAGGCTTGTGTTTGTGTCCTGAAAAATGATACGCGAACTGTAATGATACGCGAACTGT |
| *int* | GCGATCAAGCCATGAATGTAACGTAACGGA | TGATACGCGAACTGTAATGATACGCGAACTGTAAGCGATCAAGCCATGAATGTAACGTAACGGAAATGATACGCGAACTGTAATGATACGCGAACTGT |
| *int* | CATCAGGCAGACCCCTCCTTATTGCTTTAA | TGATACGCGAACTGTAATGATACGCGAACTGTAACATCAGGCAGACCCCTCCTTATTGCTTTAAAATGATACGCGAACTGTAATGATACGCGAACTGT |
| *int* | CGTCTATGTATCCATTGAGCATTGCCGCAA | TGATACGCGAACTGTAATGATACGCGAACTGTAACGTCTATGTATCCATTGAGCATTGCCGCAAAATGATACGCGAACTGTAATGATACGCGAACTGT |
| *int* | TGCCTCTCGGAATGCATCGCTCAGTGTTGA | TGATACGCGAACTGTAATGATACGCGAACTGTAATGCCTCTCGGAATGCATCGCTCAGTGTTGAAATGATACGCGAACTGTAATGATACGCGAACTGT |
| *int* | TCTCCTTACCTCTGATTTTGCTGCGCGAGT | TGATACGCGAACTGTAATGATACGCGAACTGTAATCTCCTTACCTCTGATTTTGCTGCGCGAGTAATGATACGCGAACTGTAATGATACGCGAACTGT |
| *int* | GCCAACATGGTGATGATTCTGCTGCTTGAT | TGATACGCGAACTGTAATGATACGCGAACTGTAAGCCAACATGGTGATGATTCTGCTGCTTGATAATGATACGCGAACTGTAATGATACGCGAACTGT |
| *int* | CATTTCGCATAAATCACCAACTCGTTGCCC | TGATACGCGAACTGTAATGATACGCGAACTGTAACATTTCGCATAAATCACCAACTCGTTGCCCAATGATACGCGAACTGTAATGATACGCGAACTGT |
| *int* | GCATCAATATGCAATGCTGTTGGGATGGCA | TGATACGCGAACTGTAATGATACGCGAACTGTAAGCATCAATATGCAATGCTGTTGGGATGGCAAATGATACGCGAACTGTAATGATACGCGAACTGT |
| *int* | CGAGTAGATGCAATTATGGTTTCTCCGCCA | TGATACGCGAACTGTAATGATACGCGAACTGTAACGAGTAGATGCAATTATGGTTTCTCCGCCAAATGATACGCGAACTGTAATGATACGCGAACTGT |
| *int* | CTGATGCTTTTCGTGCGCGCATAAAATACC | TGATACGCGAACTGTAATGATACGCGAACTGTAACTGATGCTTTTCGTGCGCGCATAAAATACCAATGATACGCGAACTGTAATGATACGCGAACTGT |
| *int* | GAGTCTTGCAGACAAACTGCGCAACTCGTG | TGATACGCGAACTGTAATGATACGCGAACTGTAAGAGTCTTGCAGACAAACTGCGCAACTCGTGAATGATACGCGAACTGTAATGATACGCGAACTGT |
| *int* | CCCGAGAAGATGTTGAGCAAACTTATCGCT | TGATACGCGAACTGTAATGATACGCGAACTGTAACCCGAGAAGATGTTGAGCAAACTTATCGCTAATGATACGCGAACTGTAATGATACGCGAACTGT |
| *int* | ATTTTGTCCCACTCCCTGCCTCTGTCATCA | TGATACGCGAACTGTAATGATACGCGAACTGTAAATTTTGTCCCACTCCCTGCCTCTGTCATCAAATGATACGCGAACTGTAATGATACGCGAACTGT |
| *gam* | GTTCATGGCTGAACTCCTGAAATAGCTGTG | GGGTGGAATTTCGTAAAGGGTGGAATTTCGTAAAGTTCATGGCTGAACTCCTGAAATAGCTGTGAAGGGTGGAATTTCGTAAAGGGTGGAATTTCGTA |
| *gam* | CCAGCTCTGAGCCTCAAGACGATCCTGAAT | GGGTGGAATTTCGTAAAGGGTGGAATTTCGTAAACCAGCTCTGAGCCTCAAGACGATCCTGAATAAGGGTGGAATTTCGTAAAGGGTGGAATTTCGTA |
| *gam* | CTCTTTCTCTTCACGGGCGAGCTGCTGGTA | GGGTGGAATTTCGTAAAGGGTGGAATTTCGTAAACTCTTTCTCTTCACGGGCGAGCTGCTGGTAAAGGGTGGAATTTCGTAAAGGGTGGAATTTCGTA |
| *gam* | TTGCTGGCCCCGTGGCGTTGCAAATGATCG | GGGTGGAATTTCGTAAAGGGTGGAATTTCGTAAATTGCTGGCCCCGTGGCGTTGCAAATGATCGAAGGGTGGAATTTCGTAAAGGGTGGAATTTCGTA |
| *gam* | AACTCAACATCGTCATCAAACGCACGGGTA | GGGTGGAATTTCGTAAAGGGTGGAATTTCGTAAAAACTCAACATCGTCATCAAACGCACGGGTAAAGGGTGGAATTTCGTAAAGGGTGGAATTTCGTA |
| *gam* | TCAACCATGTACCGGATGTGTTCTGCCATG | GGGTGGAATTTCGTAAAGGGTGGAATTTCGTAAATCAACCATGTACCGGATGTGTTCTGCCATGAAGGGTGGAATTTCGTAAAGGGTGGAATTTCGTA |
| *kil* | GTGGCGATAGTGAATTTAGTCTGGATAGCC | GGGTGGAATTTCGTAAAGGGTGGAATTTCGTAAAGTGGCGATAGTGAATTTAGTCTGGATAGCCAAGGGTGGAATTTCGTAAAGGGTGGAATTTCGTA |
| *kil* | TCGACGGCTTCACGAAACATCTTTTCATCG | GGGTGGAATTTCGTAAAGGGTGGAATTTCGTAAATCGACGGCTTCACGAAACATCTTTTCATCGAAGGGTGGAATTTCGTAAAGGGTGGAATTTCGTA |
| *cIII* | AACAGGCCACCCTGCAATGGCATATTGCAT | GGGTGGAATTTCGTAAAGGGTGGAATTTCGTAAAAACAGGCCACCCTGCAATGGCATATTGCATAAGGGTGGAATTTCGTAAAGGGTGGAATTTCGTA |
| *cIII* | CCGTCACGTAATTTACGGGTGATTCGTTCA | GGGTGGAATTTCGTAAAGGGTGGAATTTCGTAAACCGTCACGTAATTTACGGGTGATTCGTTCAAAGGGTGGAATTTCGTAAAGGGTGGAATTTCGTA |
| *cIII* | GGGACTCCTGGCTGATTAAGTATGTCGATA | GGGTGGAATTTCGTAAAGGGTGGAATTTCGTAAAGGGACTCCTGGCTGATTAAGTATGTCGATAAAGGGTGGAATTTCGTAAAGGGTGGAATTTCGTA |
| *N* | CCTTCATGGTGGTCAGTGCGTCCTGCTGAT | AGATTGGCGCGACATAAAGATTGGCGCGACATAACCTTCATGGTGGTCAGTGCGTCCTGCTGATAAAGATTGGCGCGACATAAAGATTGGCGCGACAT |
| *N* | TTGGCACATTGGCAGCTAATCCGGAATCGC | AGATTGGCGCGACATAAAGATTGGCGCGACATAATTGGCACATTGGCAGCTAATCCGGAATCGCAAAGATTGGCGCGACATAAAGATTGGCGCGACAT |
| *N* | TGTGTGGCAGTTGTAGTCCTGAACGAAAAC | AGATTGGCGCGACATAAAGATTGGCGCGACATAATGTGTGGCAGTTGTAGTCCTGAACGAAAACAAAGATTGGCGCGACATAAAGATTGGCGCGACAT |
| *N* | TGCATCCATCTGGATTCTCCTGTCAGTTAG | AGATTGGCGCGACATAAAGATTGGCGCGACATAATGCATCCATCTGGATTCTCCTGTCAGTTAGAAAGATTGGCGCGACATAAAGATTGGCGCGACAT |
| *N* | TGAGCCTGTTTCTCTGCGCGACGTTCGCGG | AGATTGGCGCGACATAAAGATTGGCGCGACATAATGAGCCTGTTTCTCTGCGCGACGTTCGCGGAAAGATTGGCGCGACATAAAGATTGGCGCGACAT |
| *N* | GCGCTTACCCCAACCAACAGGGGATTTGCT | AGATTGGCGCGACATAAAGATTGGCGCGACATAAGCGCTTACCCCAACCAACAGGGGATTTGCTAAAGATTGGCGCGACATAAAGATTGGCGCGACAT |
| *N* | TTGCGATTCAGCGAGAGAATAGGGCGGTTA | AGATTGGCGCGACATAAAGATTGGCGCGACATAATTGCGATTCAGCGAGAGAATAGGGCGGTTAAAAGATTGGCGCGACATAAAGATTGGCGCGACAT |
| *N* | GGCTTGCTGTACCATGTGCGCTGATTCTTG | AGATTGGCGCGACATAAAGATTGGCGCGACATAAGGCTTGCTGTACCATGTGCGCTGATTCTTGAAAGATTGGCGCGACATAAAGATTGGCGCGACAT |
| *N* | ATTTTCTGGCGTCCACTGCATGTTATGCCG | AGATTGGCGCGACATAAAGATTGGCGCGACATAAATTTTCTGGCGTCCACTGCATGTTATGCCGAAAGATTGGCGCGACATAAAGATTGGCGCGACAT |
| *N* | CTGGACTGAATTAGTTGCCAGCTATGATCC | AGATTGGCGCGACATAAAGATTGGCGCGACATAACTGGACTGAATTAGTTGCCAGCTATGATCCAAAGATTGGCGCGACATAAAGATTGGCGCGACAT |
| *N* | GCCCTTAATGAAGGCAGGAAGTATGTGGTT | AGATTGGCGCGACATAAAGATTGGCGCGACATAAGCCCTTAATGAAGGCAGGAAGTATGTGGTTAAAGATTGGCGCGACATAAAGATTGGCGCGACAT |
| *N* | GCCAGCGCCAGATATAAGCGATTTAAGCTA | AGATTGGCGCGACATAAAGATTGGCGCGACATAAGCCAGCGCCAGATATAAGCGATTTAAGCTAAAAGATTGGCGCGACATAAAGATTGGCGCGACAT |
| *N* | CTGCTTTTAAGCCCAGATAACTGGCCTGAA | AGATTGGCGCGACATAAAGATTGGCGCGACATAACTGCTTTTAAGCCCAGATAACTGGCCTGAAAAAGATTGGCGCGACATAAAGATTGGCGCGACAT |
| *rexB* | GTAAACACCAGGCATGATTCTGTTCCGCAT | GGGCCAGCGCAATAAAAGGGCCAGCGCAATAAAAGTAAACACCAGGCATGATTCTGTTCCGCATAAGGGCCAGCGCAATAAAAGGGCCAGCGCAATAA |
| *rexB* | GCAATCAGGATTGCAATCATGGTTCCTGCA | GGGCCAGCGCAATAAAAGGGCCAGCGCAATAAAAGCAATCAGGATTGCAATCATGGTTCCTGCAAAGGGCCAGCGCAATAAAAGGGCCAGCGCAATAA |
| *rexB* | AGTCGGCGCGTTCTGCTTCCGATTAGAAAC | GGGCCAGCGCAATAAAAGGGCCAGCGCAATAAAAAGTCGGCGCGTTCTGCTTCCGATTAGAAACAAGGGCCAGCGCAATAAAAGGGCCAGCGCAATAA |
| *rexB* | GAGGCAATGCCGATGGCGATAGTGGGTATC | GGGCCAGCGCAATAAAAGGGCCAGCGCAATAAAAGAGGCAATGCCGATGGCGATAGTGGGTATCAAGGGCCAGCGCAATAAAAGGGCCAGCGCAATAA |
| *rexA* | GCCAGCAGAGAATTAAGGAAAACAGACAGG | GGGCCAGCGCAATAAAAGGGCCAGCGCAATAAAAGCCAGCAGAGAATTAAGGAAAACAGACAGGAAGGGCCAGCGCAATAAAAGGGCCAGCGCAATAA |
| *rexA* | CAATGATTCGTCATCTGCGAGGCTGTTCTT | GGGCCAGCGCAATAAAAGGGCCAGCGCAATAAAACAATGATTCGTCATCTGCGAGGCTGTTCTTAAGGGCCAGCGCAATAAAAGGGCCAGCGCAATAA |
| *rexA* | TCTGTCAGATCGGATGTGGTCGGCCCGAAA | GGGCCAGCGCAATAAAAGGGCCAGCGCAATAAAATCTGTCAGATCGGATGTGGTCGGCCCGAAAAAGGGCCAGCGCAATAAAAGGGCCAGCGCAATAA |
| *rexA* | TCTCCACTGCTTAATGACATTCCTTTCCCG | GGGCCAGCGCAATAAAAGGGCCAGCGCAATAAAATCTCCACTGCTTAATGACATTCCTTTCCCGAAGGGCCAGCGCAATAAAAGGGCCAGCGCAATAA |
| *rexA* | TTGGTGGTTCCCCTCATCAGTGGCTCTATC | GGGCCAGCGCAATAAAAGGGCCAGCGCAATAAAATTGGTGGTTCCCCTCATCAGTGGCTCTATCAAGGGCCAGCGCAATAAAAGGGCCAGCGCAATAA |
| *rexA* | CGTTGTTCGGCCGATGAAATGCATATGCAT | GGGCCAGCGCAATAAAAGGGCCAGCGCAATAAAACGTTGTTCGGCCGATGAAATGCATATGCATAAGGGCCAGCGCAATAAAAGGGCCAGCGCAATAA |
| *rexA* | CCCTTCAATATCTGTTGCCCCTAAGACCTT | GGGCCAGCGCAATAAAAGGGCCAGCGCAATAAAACCCTTCAATATCTGTTGCCCCTAAGACCTTAAGGGCCAGCGCAATAAAAGGGCCAGCGCAATAA |
| *rexA* | TTGTTCAGAGGCGCTGAGAGATGGCCTTTT | GGGCCAGCGCAATAAAAGGGCCAGCGCAATAAAATTGTTCAGAGGCGCTGAGAGATGGCCTTTTAAGGGCCAGCGCAATAAAAGGGCCAGCGCAATAA |
| *rexA* | GCAATATGCCATCTCTTCAGCTATCTCAGC | GGGCCAGCGCAATAAAAGGGCCAGCGCAATAAAAGCAATATGCCATCTCTTCAGCTATCTCAGCAAGGGCCAGCGCAATAAAAGGGCCAGCGCAATAA |
| *cI* | CGTCCTCAAGCTGCTCTTGTGTTAATGGTT | TGCGTACGACAATGTAATGCGTACGACAATGTAACGTCCTCAAGCTGCTCTTGTGTTAATGGTTAATGCGTACGACAATGTAATGCGTACGACAATGT |
| *cI* | CGACAGATTCCTGGGATAAGCCAAGTTCAT | TGCGTACGACAATGTAATGCGTACGACAATGTAACGACAGATTCCTGGGATAAGCCAAGTTCATAATGCGTACGACAATGTAATGCGTACGACAATGT |
| *cI* | CACCAACGCCTGACTGCCCCATCCCCATCT | TGCGTACGACAATGTAATGCGTACGACAATGTAACACCAACGCCTGACTGCCCCATCCCCATCTAATGCGTACGACAATGTAATGCGTACGACAATGT |
| *cI* | GGCGATTGAAGGGCTAAATTCTTCAACGCT | TGCGTACGACAATGTAATGCGTACGACAATGTAAGGCGATTGAAGGGCTAAATTCTTCAACGCTAATGCGTACGACAATGTAATGCGTACGACAATGT |
| *cI* | GCTGCATACTAACCGCTTCATACATCTCGT | TGCGTACGACAATGTAATGCGTACGACAATGTAAGCTGCATACTAACCGCTTCATACATCTCGTAATGCGTACGACAATGTAATGCGTACGACAATGT |
| *cI* | GTGAGAACATCCCTGCCTGAACATGAGAAA | TGCGTACGACAATGTAATGCGTACGACAATGTAAGTGAGAACATCCCTGCCTGAACATGAGAAAAATGCGTACGACAATGTAATGCGTACGACAATGT |
| *cI* | CTCTCCGCATCACCTTTGGTAAAGGTTCTA | TGCGTACGACAATGTAATGCGTACGACAATGTAACTCTCCGCATCACCTTTGGTAAAGGTTCTAAATGCGTACGACAATGTAATGCGTACGACAATGT |
| *cI* | CTCAAGCCAGAATGCAGAATCACTGGCTTT | TGCGTACGACAATGTAATGCGTACGACAATGTAACTCAAGCCAGAATGCAGAATCACTGGCTTTAATGCGTACGACAATGTAATGCGTACGACAATGT |
| *cI* | GGAGCCTGTTGGTGCGGTCATGGAATTACC | TGCGTACGACAATGTAATGCGTACGACAATGTAAGGAGCCTGTTGGTGCGGTCATGGAATTACCAATGCGTACGACAATGTAATGCGTACGACAATGT |
| *cI* | GGAGCCTGTTGGTGCGGTCATGGAATTACC | TGCGTACGACAATGTAATGCGTACGACAATGTAAGGAGCCTGTTGGTGCGGTCATGGAATTACCAATGCGTACGACAATGTAATGCGTACGACAATGT |
| *cI* | GCTATGCAGAAATCACCTGGCTCAACAGCC | TGCGTACGACAATGTAATGCGTACGACAATGTAAGCTATGCAGAAATCACCTGGCTCAACAGCCAATGCGTACGACAATGTAATGCGTACGACAATGT |
| *cI* | CCGCTATCCCTGATCAGTTTCTTGAAGGTA | TGCGTACGACAATGTAATGCGTACGACAATGTAACCGCTATCCCTGATCAGTTTCTTGAAGGTAAATGCGTACGACAATGTAATGCGTACGACAATGT |
| *cI* | GATCATTGGGTACTGTGGGTTTAGTGGTTG | TGCGTACGACAATGTAATGCGTACGACAATGTAAGATCATTGGGTACTGTGGGTTTAGTGGTTGAATGCGTACGACAATGTAATGCGTACGACAATGT |
| *cI* | GATAACTTTCCCCACAACGGAACAACTCTC | TGCGTACGACAATGTAATGCGTACGACAATGTAAGATAACTTTCCCCACAACGGAACAACTCTCAATGCGTACGACAATGTAATGCGTACGACAATGT |
| *cro* | GGGTTATGCGTTGTTCCATACAACCTCCTT | GTCCCGGCGTATCAAAAGTCCCGGCGTATCAAAAGGGTTATGCGTTGTTCCATACAACCTCCTTAAGTCCCGGCGTATCAAAAGTCCCGGCGTATCAA |
| *cro* | CTTGGTTTGCCCAAAGCGCATTGCATAATC | GTCCCGGCGTATCAAAAGTCCCGGCGTATCAAAACTTGGTTTGCCCAAAGCGCATTGCATAATCAAGTCCCGGCGTATCAAAAGTCCCGGCGTATCAA |
| *cro* | CGCTTTGATATACGCCGAGATCTTTAGCTG | GTCCCGGCGTATCAAAAGTCCCGGCGTATCAAAACGCTTTGATATACGCCGAGATCTTTAGCTGAAGTCCCGGCGTATCAAAAGTCCCGGCGTATCAA |
| *cro* | TTTCGGCCTGCATGAATGGCCTTGTTGATC | GTCCCGGCGTATCAAAAGTCCCGGCGTATCAAAATTTCGGCCTGCATGAATGGCCTTGTTGATCAAGTCCCGGCGTATCAAAAGTCCCGGCGTATCAA |
| *cro* | CCGCATAAACGCTTCCATCAGCGTTTATAG | GTCCCGGCGTATCAAAAGTCCCGGCGTATCAAAACCGCATAAACGCTTCCATCAGCGTTTATAGAAGTCCCGGCGTATCAAAAGTCCCGGCGTATCAA |
| *cro* | GGCTGGAATGTGTAAGAGCGGGGTTATTTA | GTCCCGGCGTATCAAAAGTCCCGGCGTATCAAAAGGCTGGAATGTGTAAGAGCGGGGTTATTTAAAGTCCCGGCGTATCAAAAGTCCCGGCGTATCAA |
| *cII* | AGCCTCGTTGCGTTTGTTTGCACGAACCAT | GCAGGGTGAACCTACAAGCAGGGTGAACCTACAAAGCCTCGTTGCGTTTGTTTGCACGAACCATAAGCAGGGTGAACCTACAAGCAGGGTGAACCTAC |
| *cII* | GCTGTCTTCTCAGTTCCAAGCATTGCGATT | GCAGGGTGAACCTACAAGCAGGGTGAACCTACAAGCTGTCTTCTCAGTTCCAAGCATTGCGATTAAGCAGGGTGAACCTACAAGCAGGGTGAACCTAC |
| *cII* | GATCTGCGACTTATCAACGCCCACAGCTTC | GCAGGGTGAACCTACAAGCAGGGTGAACCTACAAGATCTGCGACTTATCAACGCCCACAGCTTCAAGCAGGGTGAACCTACAAGCAGGGTGAACCTAC |
| *cII* | ACTTTGGAATCCAGTCCCTCTTCCACCTGC | GCAGGGTGAACCTACAAGCAGGGTGAACCTACAAACTTTGGAATCCAGTCCCTCTTCCACCTGCAAGCAGGGTGAACCTACAAGCAGGGTGAACCTAC |
| *cII* | CCCCATTCAAGAACAGCAAGCAGCATTGAG | GCAGGGTGAACCTACAAGCAGGGTGAACCTACAACCCCATTCAAGAACAGCAAGCAGCATTGAGAAGCAGGGTGAACCTACAAGCAGGGTGAACCTAC |
| *cII* | CGCCAATCGAGCCATGTCGTCGTCAACGAC | GCAGGGTGAACCTACAAGCAGGGTGAACCTACAACGCCAATCGAGCCATGTCGTCGTCAACGACAAGCAGGGTGAACCTACAAGCAGGGTGAACCTAC |
| *cII* | TCAGAACGCTCGGTTGCCGCCGGGCGTTTT | GCAGGGTGAACCTACAAGCAGGGTGAACCTACAATCAGAACGCTCGGTTGCCGCCGGGCGTTTTAAGCAGGGTGAACCTACAAGCAGGGTGAACCTAC |

### Table S4. Table S4 (continued)

| Gene | mRNA binding region sequence | Full primary probe sequence |
|---|---|---|
| *O* | GGCAAAGTTACCTCTGCCGAAGTTGAGTAT | TCTGTAACCGCAAGTAATCTGTAACCGCAAGTAAGGCAAAGTTACCTCTGCCGAAGTTGAGTATAATCTGTAACCGCAAGTAATCTGTAACCGCAAGT |
| *O* | CCAAACATGCCGCCTTGCTGCTTGATAATA | TCTGTAACCGCAAGTAATCTGTAACCGCAAGTAACCAAACATGCCGCCTTGCTGCTTGATAATAAATCTGTAACCGCAAGTAATCTGTAACCGCAAGT |
| *O* | CAGAGGATTCGCCAGAATTCTCTGACGAAT | TCTGTAACCGCAAGTAATCTGTAACCGCAAGTAACAGAGGATTCGCCAGAATTCTCTGACGAATAATCTGTAACCGCAAGTAATCTGTAACCGCAAGT |
| *O* | GGCAAGCAGCACTTTAAACTGTCGCTTGGT | TCTGTAACCGCAAGTAATCTGTAACCGCAAGTAAGGCAAGCAGCACTTTAAACTGTCGCTTGGTAATCTGTAACCGCAAGTAATCTGTAACCGCAAGT |
| *O* | TGGCTGATGGTGCGATAGTCTTCACCATGT | TCTGTAACCGCAAGTAATCTGTAACCGCAAGTAATGGCTGATGGTGCGATAGTCTTCACCATGTAATCTGTAACCGCAAGTAATCTGTAACCGCAAGT |
| *O* | ACATGTCGCGGTGGTTACGTCCGTCACGTT | TCTGTAACCGCAAGTAATCTGTAACCGCAAGTAAACATGTCGCGGTGGTTACGTCCGTCACGTTAATCTGTAACCGCAAGTAATCTGTAACCGCAAGT |
| *O* | ACCAGAAGTTGTCCTGGCATGCCCAGCGGA | TCTGTAACCGCAAGTAATCTGTAACCGCAAGTAAACCAGAAGTTGTCCTGGCATGCCCAGCGGAAATCTGTAACCGCAAGTAATCTGTAACCGCAAGT |
| *O* | ACTTATCGCGGAGTTTGGCCGGGCTCAGCA | TCTGTAACCGCAAGTAATCTGTAACCGCAAGTAAACTTATCGCGGAGTTTGGCCGGGCTCAGCAAATCTGTAACCGCAAGTAATCTGTAACCGCAAGT |
| *P* | TTCCGGCATGTTGTTGGCGATCCGACGCAT | TCTGTAACCGCAAGTAATCTGTAACCGCAAGTAATTCCGGCATGTTGTTGGCGATCCGACGCATAATCTGTAACCGCAAGTAATCTGTAACCGCAAGT |
| *P* | GAAAGTTGCCAGTAACTGGCTGAACACACC | TCTGTAACCGCAAGTAATCTGTAACCGCAAGTAAGAAAGTTGCCAGTAACTGGCTGAACACACCAATCTGTAACCGCAAGTAATCTGTAACCGCAAGT |
| *P* | GTTCACTTCGTTCTGGTCACGGTTAGCCAG | TCTGTAACCGCAAGTAATCTGTAACCGCAAGTAAGTTCACTTCGTTCTGGTCACGGTTAGCCAGAATCTGTAACCGCAAGTAATCTGTAACCGCAAGT |
| *P* | GGGTGATGGCAGAAATGGTCGATTCTGCCG | TCTGTAACCGCAAGTAATCTGTAACCGCAAGTAAGGGTGATGGCAGAAATGGTCGATTCTGCCGAATCTGTAACCGCAAGTAATCTGTAACCGCAAGT |
| *P* | CGCATTGGCCCGCATGTTCTGATACAGGTT | TCTGTAACCGCAAGTAATCTGTAACCGCAAGTAACGCATTGGCCCGCATGTTCTGATACAGGTTAATCTGTAACCGCAAGTAATCTGTAACCGCAAGT |
| *P* | CGCCTCACCACGGTTAATTCTCGCAGTCAT | TCTGTAACCGCAAGTAATCTGTAACCGCAAGTAACGCCTCACCACGGTTAATTCTCGCAGTCATAATCTGTAACCGCAAGTAATCTGTAACCGCAAGT |
| *P* | TGCGATCTTCGCCAGAGCCTGTGCACGATT | TCTGTAACCGCAAGTAATCTGTAACCGCAAGTAATGCGATCTTCGCCAGAGCCTGTGCACGATTAATCTGTAACCGCAAGTAATCTGTAACCGCAAGT |
| *Q* | CCGTGGTGAGTCGCTCATCATCGGGCTTTT | GCGCAATAAACCCTAAAGCGCAATAAACCCTAAACCGTGGTGAGTCGCTCATCATCGGGCTTTTAAGCGCAATAAACCCTAAAGCGCAATAAACCCTA |
| *Q* | CAGTACCGGAAAGAGAGTCAGAAGCCGTGG | GCGCAATAAACCCTAAAGCGCAATAAACCCTAAACAGTACCGGAAAGAGAGTCAGAAGCCGTGGAAGCGCAATAAACCCTAAAGCGCAATAAACCCTA |
| *Q* | GATTGCGCCATCCCCATAGCAGCCATCACA | GCGCAATAAACCCTAAAGCGCAATAAACCCTAAAGATTGCGCCATCCCCATAGCAGCCATCACAAAGCGCAATAAACCCTAAAGCGCAATAAACCCTA |
| *Q* | GCAGAATGCAGCCATACCGAATCCGGCTTG | GCGCAATAAACCCTAAAGCGCAATAAACCCTAAAGCAGAATGCAGCCATACCGAATCCGGCTTGAAGCGCAATAAACCCTAAAGCGCAATAAACCCTA |
| *Q* | GTTTGTCGTTCTGGCTGAGTTCGTGCTTAC | GCGCAATAAACCCTAAAGCGCAATAAACCCTAAAGTTTGTCGTTCTGGCTGAGTTCGTGCTTACAAGCGCAATAAACCCTAAAGCGCAATAAACCCTA |
| *Q* | CCCGATACCTTGTGTGCAAATTGCATCAGA | GCGCAATAAACCCTAAAGCGCAATAAACCCTAAACCCGATACCTTGTGTGCAAATTGCATCAGAAAGCGCAATAAACCCTAAAGCGCAATAAACCCTA |
| *Q* | TCCTTCAAGCTTTGCCACACCACGGTATTT | GCGCAATAAACCCTAAAGCGCAATAAACCCTAAATCCTTCAAGCTTTGCCACACCACGGTATTTAAGCGCAATAAACCCTAAAGCGCAATAAACCCTA |
| *Q* | CGAGCACTTGCAGTACCTTTGCCTTAGTAT | GCGCAATAAACCCTAAAGCGCAATAAACCCTAAACGAGCACTTGCAGTACCTTTGCCTTAGTATAAGCGCAATAAACCCTAAAGCGCAATAAACCCTA |
| *Q* | CACTACGGCAATAATCCGCATAAGCGAATG | GCGCAATAAACCCTAAAGCGCAATAAACCCTAAACACTACGGCAATAATCCGCATAAGCGAATGAAGCGCAATAAACCCTAAAGCGCAATAAACCCTA |
| *Q* | TGTACCATGGCAATCTCTGCATCTTGCCCC | GCGCAATAAACCCTAAAGCGCAATAAACCCTAAATGTACCATGGCAATCTCTGCATCTTGCCCCAAGCGCAATAAACCCTAAAGCGCAATAAACCCTA |
| *Q* | GCTCTGTTTTGGCAATATCAACCGCACGGC | GCGCAATAAACCCTAAAGCGCAATAAACCCTAAAGCTCTGTTTTGGCAATATCAACCGCACGGCAAGCGCAATAAACCCTAAAGCGCAATAAACCCTA |
| *Q* | CTTCCGCACTCTTTCTCGACAACTCTCCCC | GCGCAATAAACCCTAAAGCGCAATAAACCCTAAACTTCCGCACTCTTTCTCGACAACTCTCCCCAAGCGCAATAAACCCTAAAGCGCAATAAACCCTA |
| *Q* | TGGCATCCTTGAATAGCCGACGCCTTTGCA | GCGCAATAAACCCTAAAGCGCAATAAACCCTAAATGGCATCCTTGAATAGCCGACGCCTTTGCAAAGCGCAATAAACCCTAAAGCGCAATAAACCCTA |
| *Q* | GCATCGTCACAGCGCGATATGCTGCGCTTG | GCGCAATAAACCCTAAAGCGCAATAAACCCTAAAGCATCGTCACAGCGCGATATGCTGCGCTTGAAGCGCAATAAACCCTAAAGCGCAATAAACCCTA |
| *Q* | GACCAGGTGGGTTGGGTAAGGTTTGGGATT | GCGCAATAAACCCTAAAGCGCAATAAACCCTAAAGACCAGGTGGGTTGGGTAAGGTTTGGGATTAAGCGCAATAAACCCTAAAGCGCAATAAACCCTA |
| *Q* | CAGAGCGTCATACAGCGGCTTAACAGTGCG | GCGCAATAAACCCTAAAGCGCAATAAACCCTAAACAGAGCGTCATACAGCGGCTTAACAGTGCGAAGCGCAATAAACCCTAAAGCGCAATAAACCCTA |
| *Q* | CGATTGACTCTTCTTTGTGGCATTGCACCA | GCGCAATAAACCCTAAAGCGCAATAAACCCTAAACGATTGACTCTTCTTTGTGGCATTGCACCAAAGCGCAATAAACCCTAAAGCGCAATAAACCCTA |
| *R* | CAGCATATCGAGGAACGCCTTACGTTGATT | CGCGCAGACGTTAATAACGCGCAGACGTTAATAACAGCATATCGAGGAACGCCTTACGTTGATTAACGCGCAGACGTTAATAACGCGCAGACGTTAAT |
| *R* | GACGTCCGTTATCAGTTCCCTCCGACCACG | CGCGCAGACGTTAATAACGCGCAGACGTTAATAAGACGTCCGTTATCAGTTCCCTCCGACCACGAACGCGCAGACGTTAATAACGCGCAGACGTTAAT |
| *R* | CTCCGCCTACAATGACGTCATAACCATGAT | CGCGCAGACGTTAATAACGCGCAGACGTTAATAACTCCGCCTACAATGACGTCATAACCATGATAACGCGCAGACGTTAATAACGCGCAGACGTTAAT |
| *R* | CGAGGGTGATCGGAGTAATCAGTAAATAGC | CGCGCAGACGTTAATAACGCGCAGACGTTAATAACGAGGGTGATCGGAGTAATCAGTAAATAGCAACGCGCAGACGTTAATAACGCGCAGACGTTAAT |
| *R* | CGCCTGTTGATTTGAGTTTTGGGTTTAGCG | CGCGCAGACGTTAATAACGCGCAGACGTTAATAACGCCTGTTGATTTGAGTTTTGGGTTTAGCGAACGCGCAGACGTTAATAACGCGCAGACGTTAAT |
| *R* | CACCAACGGGAAAGAAGCTGGTAGCGTCCG | CGCGCAGACGTTAATAACGCGCAGACGTTAATAACACCAACGGGAAAGAAGCTGGTAGCGTCCGAACGCGCAGACGTTAATAACGCGCAGACGTTAAT |
| *R* | TTTCAGGCCAAGCTGCTTGCGGTAGGCATC | CGCGCAGACGTTAATAACGCGCAGACGTTAATAATTTCAGGCCAAGCTGCTTGCGGTAGGCATCAACGCGCAGACGTTAATAACGCGCAGACGTTAAT |
| *R* | CCACAGCGTCCTGACTTTTCGGAGAGAAGT | CGCGCAGACGTTAATAACGCGCAGACGTTAATAACCACAGCGTCCTGACTTTTCGGAGAGAAGTAACGCGCAGACGTTAATAACGCGCAGACGTTAAT |
| *R* | AAAGCGCCACGCTCCTTAATCTGCTGCAAT | CGCGCAGACGTTAATAACGCGCAGACGTTAATAAAAAGCGCCACGCTCCTTAATCTGCTGCAATAACGCGCAGACGTTAATAACGCGCAGACGTTAAT |
| *R* | CTGACGGATATCACCACGATCAATCATAGG | CGCGCAGACGTTAATAACGCGCAGACGTTAATAACTGACGGATATCACCACGATCAATCATAGGAACGCGCAGACGTTAATAACGCGCAGACGTTAAT |
| *R* | AAGCCCAGATATTGCTGCAACGGTCGATTG | CGCGCAGACGTTAATAACGCGCAGACGTTAATAAAAGCCCAGATATTGCTGCAACGGTCGATTGAACGCGCAGACGTTAATAACGCGCAGACGTTAAT |
| *R* | TCGAACTGACCATAACCAGCGCCCGGCAGT | CGCGCAGACGTTAATAACGCGCAGACGTTAATAATCGAACTGACCATAACCAGCGCCCGGCAGTAACGCGCAGACGTTAATAACGCGCAGACGTTAAT |
| *R* | CCCGCTTCTTTGAATTTTGCAATCAGGCTG | CGCGCAGACGTTAATAACGCGCAGACGTTAATAACCCGCTTCTTTGAATTTTGCAATCAGGCTGAACGCGCAGACGTTAATAACGCGCAGACGTTAAT |
| *A* | CGCATACCAGGAAGGGCGCTGGGAAACACT | TGCTCCCGGATTACAAATGCTCCCGGATTACAAAAGTGTTTCCCAGCGCCCTTCCTGGTATGCGAATGCTCCCGGATTACAAATGCTCCCGGATTACA |
| *A* | ACACGGGCAGACTTCACCACATTCACCTCA | TGCTCCCGGATTACAAATGCTCCCGGATTACAAAACACGGGCAGACTTCACCACATTCACCTCAAATGCTCCCGGATTACAAATGCTCCCGGATTACA |
| *A* | GTTCTCGGCATCACCATCCGTCGGCAACCA | TGCTCCCGGATTACAAATGCTCCCGGATTACAAAGTTCTCGGCATCACCATCCGTCGGCAACCAAATGCTCCCGGATTACAAATGCTCCCGGATTACA |
| *A* | TCATGGTGAGCGTGTTATCCCGGTGCTTTT | TGCTCCCGGATTACAAATGCTCCCGGATTACAAATCATGGTGAGCGTGTTATCCCGGTGCTTTTAATGCTCCCGGATTACAAATGCTCCCGGATTACA |
| *A* | CATAACCCGCCACATCCACCGACTTTTCAC | TGCTCCCGGATTACAAATGCTCCCGGATTACAAACATAACCCGCCACATCCACCGACTTTTCACAATGCTCCCGGATTACAAATGCTCCCGGATTACA |
| *A* | CCAGACCGAGCCTTCAATACGCTTGTCACC | TGCTCCCGGATTACAAATGCTCCCGGATTACAAACCAGACCGAGCCTTCAATACGCTTGTCACCAATGCTCCCGGATTACAAATGCTCCCGGATTACA |
| *A* | GCCTCTCACTTTTGGCGTGGAGCCACGGAT | TGCTCCCGGATTACAAATGCTCCCGGATTACAAAGCCTCTCACTTTTGGCGTGGAGCCACGGATAATGCTCCCGGATTACAAATGCTCCCGGATTACA |
| *A* | ATGCGGGGATTCACTGGCTGCACGCTCAAT | TGCTCCCGGATTACAAATGCTCCCGGATTACAAAATGCGGGGATTCACTGGCTGCACGCTCAATAATGCTCCCGGATTACAAATGCTCCCGGATTACA |
| *A* | TGACGAAAACCAGAGAATGCCATCACGGGT | TGCTCCCGGATTACAAATGCTCCCGGATTACAAATGACGAAAACCAGAGAATGCCATCACGGGTAATGCTCCCGGATTACAAATGCTCCCGGATTACA |
| *A* | CTTCAGCATCCGGACGTTCGCCAATTTTCG | TGCTCCCGGATTACAAATGCTCCCGGATTACAAACTTCAGCATCCGGACGTTCGCCAATTTTCGAATGCTCCCGGATTACAAATGCTCCCGGATTACA |
| *E* | CAGCAGTTGGGCGGTTGTGTACATCGACAT | AATGACGCGACATAGAAAATGACGCGACATAGAACAGCAGTTGGGCGGTTGTGTACATCGACATAAAATGACGCGACATAGAAAATGACGCGACATAG |
| *E* | GATAACCTCACCGGAAACAATCGGCGAAAC | AATGACGCGACATAGAAAATGACGCGACATAGAAGATAACCTCACCGGAAACAATCGGCGAAACAAAATGACGCGACATAGAAAATGACGCGACATAG |
| *E* | GGTCATCTGCGGATTCACTTCATGCTTCGG | AATGACGCGACATAGAAAATGACGCGACATAGAAGGTCATCTGCGGATTCACTTCATGCTTCGGAAAATGACGCGACATAGAAAATGACGCGACATAG |
| *E* | CGCCAGATTCTGCGGATCTTCATCCGGCAG | AATGACGCGACATAGAAAATGACGCGACATAGAACGCCAGATTCTGCGGATCTTCATCCGGCAGAAAATGACGCGACATAGAAAATGACGCGACATAG |
| *E* | CTCTTCGACCTGAGCAATGGCCAGCTCTTC | AATGACGCGACATAGAAAATGACGCGACATAGAACTCTTCGACCTGAGCAATGGCCAGCTCTTCAAAATGACGCGACATAGAAAATGACGCGACATAG |
| *E* | CATATCCACCTCAACCGGATCGAAGGCTTC | AATGACGCGACATAGAAAATGACGCGACATAGAACATATCCACCTCAACCGGATCGAAGGCTTCAAAATGACGCGACATAGAAAATGACGCGACATAG |
| *E* | CGGGTCATACGTGGACTTGTCACGCTTGCT | AATGACGCGACATAGAAAATGACGCGACATAGAACGGGTCATACGTGGACTTGTCACGCTTGCTAAAATGACGCGACATAGAAAATGACGCGACATAG |
| *E* | CGCCCAGCCTTTCGGATCGAACACGATGAT | AATGACGCGACATAGAAAATGACGCGACATAGAACGCCCAGCCTTTCGGATCGAACACGATGATAAAATGACGCGACATAGAAAATGACGCGACATAG |
| *E* | CACCGCTGTCTCCAGCTCGGAATTAGAGCC | AATGACGCGACATAGAAAATGACGCGACATAGAACACCGCTGTCTCCAGCTCGGAATTAGAGCCAAAATGACGCGACATAGAAAATGACGCGACATAG |
| *E* | CACGTACTGTCCGGAATACACGACGATGGC | AATGACGCGACATAGAAAATGACGCGACATAGAACACGTACTGTCCGGAATACACGACGATGGCAAAATGACGCGACATAGAAAATGACGCGACATAG |
| *E* | CCATCGTGTTGTCCGGCAGGAAGTTCTTTT | AATGACGCGACATAGAAAATGACGCGACATAGAACCATCGTGTTGTCCGGCAGGAAGTTCTTTTAAAATGACGCGACATAGAAAATGACGCGACATAG |
| *E* | TAATGCCTTCGCGCTGTGCGTCCGCATCCT | AATGACGCGACATAGAAAATGACGCGACATAGAATAATGCCTTCGCGCTGTGCGTCCGCATCCTAAAATGACGCGACATAGAAAATGACGCGACATAG |
| *V* | CCAGTCAACGTCTGAAAGCGGATTCGCGTA | CTCAACGTCCATGTCAACTCAACGTCCATGTCAACCAGTCAACGTCTGAAAGCGGATTCGCGTAAACTCAACGTCCATGTCAACTCAACGTCCATGTC |
| *V* | ATCATCGAGATAGCTGTCGTCATAGGACTC | CTCAACGTCCATGTCAACTCAACGTCCATGTCAAATCATCGAGATAGCTGTCGTCATAGGACTCAACTCAACGTCCATGTCAACTCAACGTCCATGTC |
| *V* | ATTTCTGCCCCTGCCCGGTCGCAGTCCAGT | CTCAACGTCCATGTCAACTCAACGTCCATGTCAAATTTCTGCCCCTGCCCGGTCGCAGTCCAGTAACTCAACGTCCATGTCAACTCAACGTCCATGTC |
| *V* | CGGGCATCCACGCCAGCGTGAAGCTGGTAT | CTCAACGTCCATGTCAACTCAACGTCCATGTCAACGGGCATCCACGCCAGCGTGAAGCTGGTATAACTCAACGTCCATGTCAACTCAACGTCCATGTC |
| *V* | CATTAAACCACGCCAGCAGCGCCTGCTGCC | CTCAACGTCCATGTCAACTCAACGTCCATGTCAACATTAAACCACGCCAGCAGCGCCTGCTGCCAACTCAACGTCCATGTCAACTCAACGTCCATGTC |
| *V* | CGATACTGCTGACCCAGCCACGGAACACAT | CTCAACGTCCATGTCAACTCAACGTCCATGTCAACGATACTGCTGACCCAGCCACGGAACACATAACTCAACGTCCATGTCAACTCAACGTCCATGTC |
| *V* | CCGTGCGGGTGATCACTTCCTTCGCCGTCA | CTCAACGTCCATGTCAACTCAACGTCCATGTCAACCGTGCGGGTGATCACTTCCTTCGCCGTCAAACTCAACGTCCATGTCAACTCAACGTCCATGTC |
| *V* | CTTCTGCCATCGACGGACGTCCCACATTGG | CTCAACGTCCATGTCAACTCAACGTCCATGTCAACTTCTGCCATCGACGGACGTCCCACATTGGAACTCAACGTCCATGTCAACTCAACGTCCATGTC |
| *V* | TCACGGTCATGCCGGTTGCCGCTGTTACCG | CTCAACGTCCATGTCAACTCAACGTCCATGTCAATCACGGTCATGCCGGTTGCCGCTGTTACCGAACTCAACGTCCATGTCAACTCAACGTCCATGTC |
| *V* | TCTTGTCGGTTACGCCCTCCGGCTGGAAGG | CTCAACGTCCATGTCAACTCAACGTCCATGTCAATCTTGTCGGTTACGCCCTCCGGCTGGAAGGAACTCAACGTCCATGTCAACTCAACGTCCATGTC |
| *V* | CAACGCCGTTCACGGTGATGGTCATACCAC | CTCAACGTCCATGTCAACTCAACGTCCATGTCAACAACGCCGTTCACGGTGATGGTCATACCACAACTCAACGTCCATGTCAACTCAACGTCCATGTC |
| *V* | TGGCGGTGACGGTAATTTCTGCAACCGCAG | CTCAACGTCCATGTCAACTCAACGTCCATGTCAATGGCGGTGACGGTAATTTCTGCAACCGCAGAACTCAACGTCCATGTCAACTCAACGTCCATGTC |

#### Table S4. Table S4 (continued)

| Gene | mRNA binding region sequence | Full primary probe sequence |
|---|---|---|
| *J* | CAACTGCGTGGACTTCAGGTTGTCCTTCGC | AGCTCGGCCTGTGATAAAGCTCGGCCTGTGATAACAACTGCGTGGACTTCAGGTTGTCCTTCGCAAAGCTCGGCCTGTGATAAAGCTCGGCCTGTGAT |
| *J* | GTGCCTGTACACCGAAGGTAAAGCGCAGAC | AGCTCGGCCTGTGATAAAGCTCGGCCTGTGATAAGTGCCTGTACACCGAAGGTAAAGCGCAGACAAAGCTCGGCCTGTGATAAAGCTCGGCCTGTGAT |
| *J* | AGCCACCGTTACGTTGTATCTGAACCAGCA | AGCTCGGCCTGTGATAAAGCTCGGCCTGTGATAAAGCCACCGTTACGTTGTATCTGAACCAGCAAAAGCTCGGCCTGTGATAAAGCTCGGCCTGTGAT |
| *J* | TACCGCTGTATTGCCGCGTCTGCGGGTTAT | AGCTCGGCCTGTGATAAAGCTCGGCCTGTGATAATACCGCTGTATTGCCGCGTCTGCGGGTTATAAAGCTCGGCCTGTGATAAAGCTCGGCCTGTGAT |
| *J* | CTGACTGGTCGCAGTACTGGCCGATGACAT | AGCTCGGCCTGTGATAAAGCTCGGCCTGTGATAACTGACTGGTCGCAGTACTGGCCGATGACATAAAGCTCGGCCTGTGATAAAGCTCGGCCTGTGAT |
| *J* | CCGGCATCACCACATTACTGCGGTTATAGG | AGCTCGGCCTGTGATAAAGCTCGGCCTGTGATAACCGGCATCACCACATTACTGCGGTTATAGGAAAGCTCGGCCTGTGATAAAGCTCGGCCTGTGAT |
| *J* | TCTTCAACAAGCTCTGTCGCCGTCTCCCAG | AGCTCGGCCTGTGATAAAGCTCGGCCTGTGATAATCTTCAACAAGCTCTGTCGCCGTCTCCCAGAAAGCTCGGCCTGTGATAAAGCTCGGCCTGTGAT |
| *J* | AATAACATCGCCCGGTACATGGCGAAGCCC | AGCTCGGCCTGTGATAAAGCTCGGCCTGTGATAAAATAACATCGCCCGGTACATGGCGAAGCCCAAAGCTCGGCCTGTGATAAAGCTCGGCCTGTGAT |
| *J* | CAGACGGAGCAGGAAACTCACGCCCTTCAC | AGCTCGGCCTGTGATAAAGCTCGGCCTGTGATAACAGACGGAGCAGGAAACTCACGCCCTTCACAAAGCTCGGCCTGTGATAAAGCTCGGCCTGTGAT |
| *J* | ATTGATACTGGCGGCTATCCAGTACAGCGC | AGCTCGGCCTGTGATAAAGCTCGGCCTGTGATAAATTGATACTGGCGGCTATCCAGTACAGCGCAAAGCTCGGCCTGTGATAAAGCTCGGCCTGTGAT |
| *J* | AACTGGCTCAGTTTGCCTTCCTCCGTGTCC | AGCTCGGCCTGTGATAAAGCTCGGCCTGTGATAAAACTGGCTCAGTTTGCCTTCCTCCGTGTCCAAAGCTCGGCCTGTGATAAAGCTCGGCCTGTGAT |
| *J* | GTCAGGCGCTTCAGGAACACGTCGTTCATG | AGCTCGGCCTGTGATAAAGCTCGGCCTGTGATAAGTCAGGCGCTTCAGGAACACGTCGTTCATGAAAGCTCGGCCTGTGATAAAGCTCGGCCTGTGAT |
| *J* | CGTTGTCCTGCCGCTGACAGTACGCTTACT | AGCTCGGCCTGTGATAAAGCTCGGCCTGTGATAACGTTGTCCTGCCGCTGACAGTACGCTTACTAAAGCTCGGCCTGTGATAAAGCTCGGCCTGTGAT |
| *J* | CACCATAACCTGCACATCGCTGGCAAACGT | AGCTCGGCCTGTGATAAAGCTCGGCCTGTGATAACACCATAACCTGCACATCGCTGGCAAACGTAAAGCTCGGCCTGTGATAAAGCTCGGCCTGTGAT |

**Table S5. 16S rRNA time-point tagging probe sequences.**

| Probe_id | Tagged sample | Full probe sequence |
|---|---|---|
| Ribo-Tag_1 | 0 min | CGATTAGTCGTCACTAA**GCGTGGACTACCAGGGTATCTAATCCTG**AAACTCCGAATGCTACG |
| Ribo-Tag_2 | 1 min | CGATTAGTCGTCACTAA**GCGTGGACTACCAGGGTATCTAATCCTG**AAGGTTACACGCGACTA |
| Ribo-Tag_3 | 2.5 min | CGATTAGTCGTCACTAA**GCGTGGACTACCAGGGTATCTAATCCTG**AATCCAGCTTACGTTCG |
| Ribo-Tag_4 | 5 min | CGATTAGTCGTCACTAA**GCGTGGACTACCAGGGTATCTAATCCTG**AAATGTAACCAAGCGTC |
| Ribo-Tag_5 | 7.5 min | CGATTAGTCGTCACTAA**GCGTGGACTACCAGGGTATCTAATCCTG**AATCAGTTACCGGTGTA |
| Ribo-Tag_6 | 10 min | ACTCCGAATGCTACGAA**GCGTGGACTACCAGGGTATCTAATCCTG**AAGGTTACACGCGACTA |
| Ribo-Tag_7 | 15 min | ACTCCGAATGCTACGAA**GCGTGGACTACCAGGGTATCTAATCCTG**AATCCAGCTTACGTTCG |
| Ribo-Tag_8 | 20 min | ATGTAACCAAGCGTCAA**GCGTGGACTACCAGGGTATCTAATCCTG**AAACTCCGAATGCTACG |
| Ribo-Tag_9 | 30 min | TCAGTTACCGGTGTAAA**GCGTGGACTACCAGGGTATCTAATCCTG**AAACTCCGAATGCTACG |
| Ribo-Tag_10 | 45 min | TCCAGCTTACGTTCGAA**GCGTGGACTACCAGGGTATCTAATCCTG**AAGGTTACACGCGACTA |
| Ribo-Tag_11 | 60 min | ATGTAACCAAGCGTCAA**GCGTGGACTACCAGGGTATCTAATCCTG**AAGGTTACACGCGACTA |
| Ribo-Tag_12 | Lysogen control | TCAGTTACCGGTGTAAA**GCGTGGACTACCAGGGTATCTAATCCTG**AAGGTTACACGCGACTA |
| Ribo-Tag_13 | Uninfected control | ATGTAACCAAGCGTCAA**GCGTGGACTACCAGGGTATCTAATCCTG**AATCCAGCTTACGTTCG |
| Ribo-Tag_14 | Not assigned | TCAGTTACCGGTGTAAA**GCGTGGACTACCAGGGTATCTAATCCTG**AATCCAGCTTACGTTCG |
| Ribo-Tag_15 | Not assigned | ATGTAACCAAGCGTCAA**GCGTGGACTACCAGGGTATCTAATCCTG**AATCAGTTACCGGTGTA |

Boldface nucleotides indicate the region targeting 16S rRNA.
